# MEMBRANE PORES ACT AS SELF-RESEALING LIPID SCRAMBLASES

**DOI:** 10.64898/2026.05.28.728368

**Authors:** Rafael B. Lira, Marco van Tilburg, Rafaela R.M. Cavalcanti, Ceri J. Richards, Karin A. Riske, Siewert J. Marrink, Wouter H. Roos

**Affiliations:** Moleculaire Biofysica, Zernike Instituut, Rijksuniversiteit Groningen. Groningen, The Netherlands; Groningen Biomolecular Sciences and Biotechnology Institute, Rijksuniversiteit Groningen. Groningen, The Netherlands; Biophysics Department, Universidade Federal de São Paulo, São Paulo, Brazil

## Abstract

Biological membranes continuously experience leaflet lipid imbalances during growth, lipid synthesis, and vesicle fusion. To alleviate such imbalances and prevent these asymmetries from compromising membrane integrity, cells rely on lipid scramblases – fast and non-specific lipid channel proteins. Here we show that lipid number asymmetry alone is sufficient to drive spontaneous formation of transient hydrophilic pores that function as self-resealing lipid scramblases. Using giant unilamellar vesicles, living cells, and coarse-grained molecular dynamics simulations, we demonstrate that fusion-induced excess lipids in one leaflet lowers membrane edge tension, generating size-selective pores whose size and lifetime scale with the magnitude of asymmetry. Below a critical threshold, these pores reseal spontaneously; above it, membranes collapse. Strikingly, pore opening enables rapid, non-selective lipid translocation between leaflets, dissipating the asymmetry that nucleates the pore and thereby promoting their own closure. Cholesterol buffers moderate imbalances through spontaneous flip flop before pore formation, whereas pore-mediated lipid scrambling relieves the remaining asymmetry and restores cholesterol’s initial distribution. Our findings identify transient lipid pores as an intrinsic, protein-independent mechanism that couples membrane destabilization to self-repair, providing a universal physical principle for membrane homeostasis during growth, remodelling, and early cellular evolution.

## Introduction

Asymmetry is ubiquitous in biology. At the cellular scale, membranes maintain strong ionic and compositional gradients between the cytoplasm and the extracellular environment, storing electrochemical and mechanical energy (1, 2). At the intracellular level, organelle identity emerges from distinct lipid and protein compositions (3). At the membrane level, lipid bilayers exhibit both lateral heterogeneity —through the formation of ordered, cholesterol- and sphingomyelin-enriched domains often termed as *rafts* (4, 5)—and transbilayer asymmetry, in which the two leaflets differ in lipid composition, packing, and elastic response (6–8). The outer leaflet of the plasma membrane (PM) is enriched in long, saturated lipids that favour dense packing and higher order, whereas the inner leaflet contains shorter, unsaturated, and charged lipids that promote fluidity (9). Due to its preference for ordered environments, cholesterol is reported to prefer the more densely packed outer leaflet (10–12). Cells invest substantial amounts of energy to create and maintain lipid compositional asymmetry/heterogeneity, underscoring its biological importance(7, 13, 14). Membrane asymmetry is thought to reconcile two competing requirements of the plasma membrane, providing a robust barrier against uncontrolled transport while preserving a fluid character (15). It has also been proposed that this asymmetry stores potential energy, which can be released when lipids are scrambled (7–9). In addition to lateral heterogeneity and transbilayer asymmetries, membranes have recently been shown to exhibit a distinct form of imbalance—lipid number asymmetry—characterized by an excess of lipids in the inner relative to the outer leaflet, with important consequences to cell function(16).

Lipid number asymmetries arise in several cellular contexts. In biogenic membranes, such as the bacterial inner membrane and the endoplasmic reticulum (ER) in eukaryotes, newly synthesized lipids are inserted predominantly into a single leaflet, reflecting the spatial localization of lipid-synthesizing enzymes(17). Additional asymmetries result from the fusion of small vesicles which, owing to their high curvature, contain more lipids in the outer than in the inner leaflet, such that their incorporation into an acceptor membrane necessarily generates leaflet imbalances. For instance, asymmetric lipid insertion in neurons produces localized membrane buckling, which functions as a mechanistic signal to initiate ultrafast membrane excision (i.e. scission)(18). Similarly, in the absence of a compensatory mechanism, the growth of synthetic cells (i.e. by lipid synthesis or vesicle fusion) is intrinsically accompanied by lipid number asymmetries(19). These imbalances translate into area mismatches, compressing the overpopulated and stretching the underpopulated leaflet. Moreover, even when the net bilayer tension is negligible, substantial stresses can be stored at the leaflet level(20, 21), with asymmetries as low as 1% being sufficient to induce membrane remodelling(22, 23).

Membranes that grow through internal lipid synthesis(24), insertion of amphiphilic molecules(25, 26), or membrane fusion(27, 28) undergo changes in both composition and leaflet area, thereby generating differential stress. Leaflet compositional asymmetry increases rigidity(29, 30), reduces edge tension (31), and a mismatch in leaflet areas makes bilayers more susceptible to rupture, as demonstrated in both simulations(32) and in experiments(33), even though it is often difficult to disentangle the changes in leaflet composition and area mismatches. Because lipids do not spontaneously translocate between membrane leaflets in intact bilayers due to the high associated energy barrier(34, 35), one pathway to relieve the ensuing stress in growing membranes is bilayer rupture via transient pore formation, which occurs once the areal strain exceeds the lysis tension(36, 37). The free energy of a circular pore of radius *r* in a lipid bilayer is

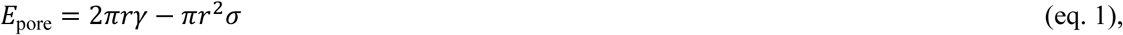

where γ is the line tension and σ the lateral membrane tension(36). At σ ∼ 0, pore nucleation is unfavourable, whereas number asymmetry generates an effective lateral tension that lowers the nucleation barrier Δ*G*^∗^ = *πγ*^2^/*σ* and defines a critical radius *r*^∗^ = *γ*/*σ*. The critical radius separates pore resealing from unstable growth(36, 38) for an effective membrane tension generated by an area difference ΔA_0_, with

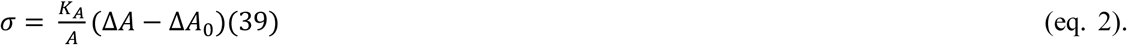

Here, K_A_ is the stretching elasticity (∼ 200 mN/m(40)) and ΔA = A_out_−A_in_ (the areas of the outer and inner leaflets, respectively). Due to high edge tension ∼ 40-50 pN(41, 42), if σ disappears, open pores in pure lipid bilayers spontaneously close in a few hundreds of milliseconds(43, 44). In contrast, at pore size > *r*\*, the pore opens indefinitely and the membrane collapses. Combining classical pore energetics (eq. 1) and area difference elasticity (eq. 2), it is possible to estimate how stress-induced asymmetries caused by changes in leaflet area modifies *r*\* via

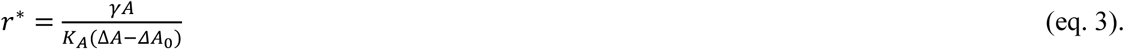

Eq. 3 shows that area imbalances (i.e. originated from lipid number asymmetries) can rupture the membrane and defines the size of the generated pore, that can either close (r < *r*\*, low-intermediate asymmetries) or lead to irreversible collapse (r > *r*\*, high asymmetries).

To prevent pore formation, leakage of essential components, and organelle collapse, biogenic membranes have evolved mechanisms to counterbalance leaflet area mismatches. Central to this regulation are proteins that catalyse lipid transport across the bilayer, such as flippases and scramblases(45). In particular, scramblases promote membrane stability during growth by mediating energy-independent, bidirectional, and non-selective phospholipid flip-flop, thereby equilibrating lipid numbers and areas between opposing leaflets(8, 46). Scramblases enable remarkably fast lipid translocation rates (∼10⁴ s⁻¹) (47), consistent with the formation of a continuous, non-selective lipid-conducting pathway across the bilayer and their proposed role as lipid channels(7).

Using a combination of reconstituted synthetic membrane, living cells, and coarse grained molecular dynamics simulations, we show that the build-up of lipid number asymmetries can compromise membrane integrity and trigger the formation of hydrophilic pores, resulting in leakage of encapsulated material, and whose size depends on the magnitude of asymmetry. When asymmetries exceed a critical threshold, *r*∗, the membrane collapses. In contrast, for moderate asymmetries, the pores that form are transient and their lifetime depends on the asymmetry. Notably, pore opening enables rapid lipid diffusion between leaflets, thereby (partially or fully) alleviating lipid number balance, and driving pore closure. This process is mechanistically analogous to scramblase activity, in which lipids translocate non-selectively, without energy input and at extremely high rates, rapidly dissipating leaflet asymmetries, making pores self-resealing structures. Cholesterol, a molecule with flipping capability, partially mitigates the initial asymmetry prior to pore formation, and pore opening subsequently re-establishes cholesterol’s original transbilayer distribution.

### Rationale

We hypothesize that the build-up of lipid-number asymmetries compromises membrane integrity; however, the structure, dynamics, fate and consequences of the resulting pores remain uncharacterized, leaving unresolved how much asymmetry synthetic and natural membranes can tolerate. To introduce defined lipid-number asymmetries, we fused large numbers of intrinsically asymmetric liposomes to synthetic membranes and living cells, and quantitatively modelled the resulting asymmetry using molecular dynamics simulations. The fusing membranes differ markedly in their physical properties: the liposomes are small, highly curved, cationic, and enriched in non–bilayer-forming lipids, giving them an intrinsic excess of outer-leaflet lipids. In contrast, the target membranes—giant unilamellar vesicles (GUVs) or HEK cells—are comparatively flat and predominantly negatively charged. GUVs and living cells also differ in their surface tension, with the former being vanishingly small, and the latter exhibiting substantially high values due to connection with the cytoskeleton. Fusion therefore changes not only membrane composition but also perturbs the physical state of the receiving membrane.

## Results

### Liposome fusion generates transient and size-selective pores in synthetic membranes

As a simplified model system, we used giant unilamellar vesicles (GUVs) that exhibit vanishingly small surface tension(48). To quantify fusion between cationic liposomes and GUVs, we developed a robust fluorescence lifetime–based FRET (FLIM-FRET) assay that reports on fusion efficiency, intermediates, and membrane mechanics, including shape changes, curvature, integrity, and pore formation. Fusion is sensitively detected at the single-GUV level by mixing of a liposomal FRET acceptor with a GUV-embedded donor (Fig. 1A). In complementary measurements, content mixing is assessed using liposomes loaded with a fluorescent reporter (Fig. 1B, i). Fusion has been shown to result in content mixing(49, 50), and thus the lack of lumenal fluorescence indicates permeabilization and leakage of the reporter (Figure 1B, ii). Because the assay is based on microscopy imaging, the associated morphological changes (e.g. membrane fluctuations, budding, or disruption) can be directly visualized, providing an additional layer of information.

**Figure 1.**
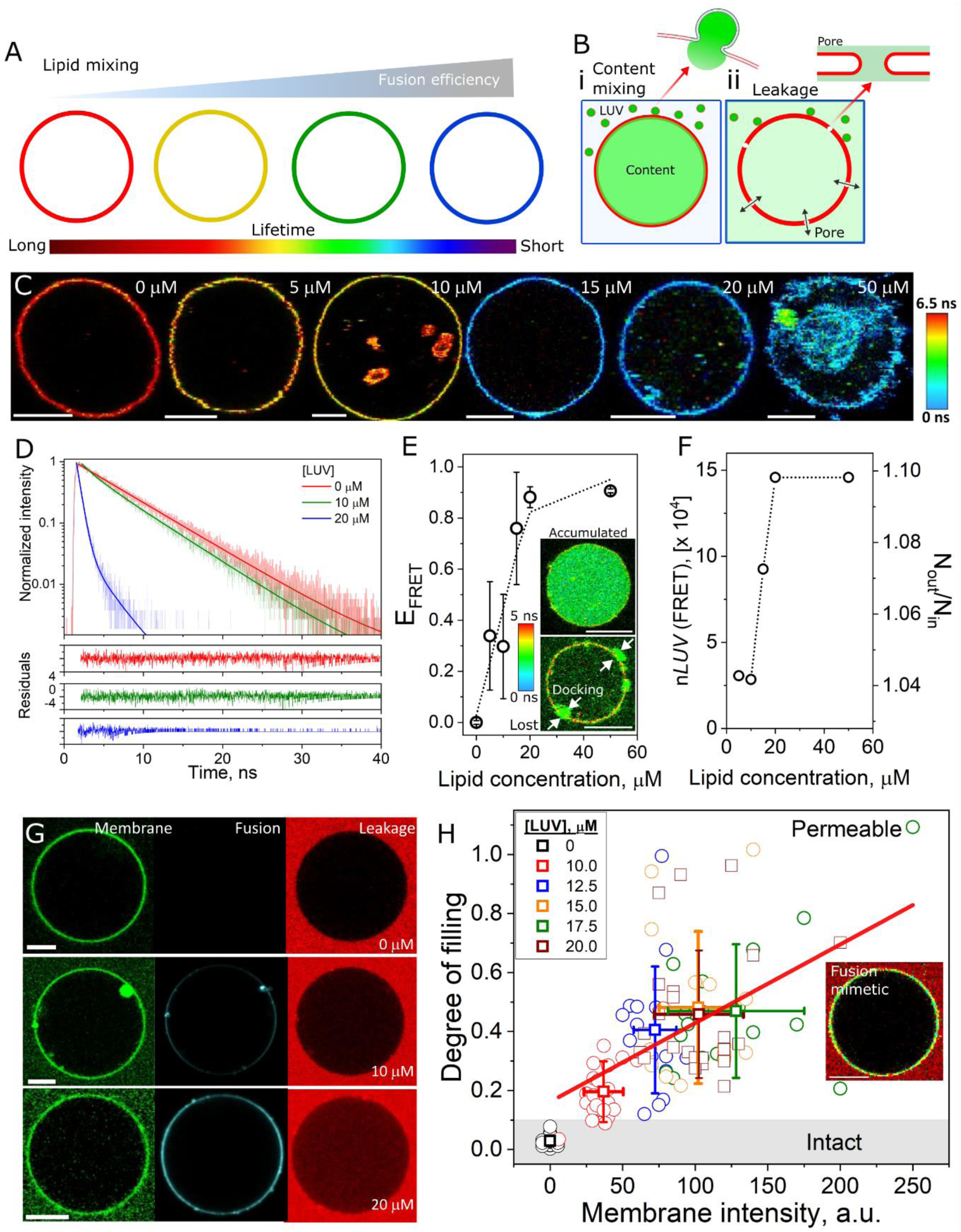
Efficient fusion of liposomes generates transient pores in GUVs. A, sketch of the fusion assay based on the transfer of a FRET acceptor lipid from fusogenic liposomes to GUVs containing a FRET donor lipid. Fusion results in an increasingly higher FRET, detected as a decrease in the fluorescence lifetime of the donor. B, sketch of the content mixing assay with liposomes encapsulating a water-soluble fluorescent probe (green) that is transferred to the GUV lumen upon full-fusion (i). If fusion is followed by pore formation, the transferred dye leaks out. C, representative images of GUVs (DOPC:_B_PS, 50:50 molar fraction - labelled with 0.5 mol% Bodipy C_16_) incubated with increasing concentrations of liposomes (DOTAP:DOPE, 50:50 molar fraction - labelled with 2 mol% DOPE-Rh). D, representative fluorescence lifetime decays measured on GUVs for various liposome concentrations. E, calculated FRET efficiency (E_FRET_) for n > 15 individual GUVs/composition for increasing concentrations of liposomes. Inset: a GUV with a clear content mixing signal of accumulated probe Dextran 10 kDa (top) and a GUV where the content has leaked out (bottom). Note the docked liposomes (arrows). F, n*LUVs* measured from FRET, and the corresponding lipid number asymmetries (N_out_/N_in_) as a function of liposome concentration. G, leakage assay on equimolar DOPC:DOPG:DOPE:Chol GUVs labelled with 0.5 mol% Bodipy C_16_ (green) incubated with increasing concentrations of DOPE-Atto647 labelled liposomes (cyan) in the presence of 5 μM SRB (red). H, correlation between the degree of filling (leakage) as a function of the membrane intensity due to fusion. Each point represents a measurement on an individual GUV. Means and errors (s.d.) are also shown. A linear fit to the data is shown (red line). The grey bar indicates the region of intact and permeable GUVs. Inset: intact GUV made of the expected lipid composition after fusion. All scale bars: 10 μm.

Increasing liposome concentration leads to a progressive decrease in donor lifetime (Figure 1 C and representative fluorescence decay curves in Figure 1D). Quantification of FRET efficiency (E_FRET,_ see methods) for n > 15 individual GUVs/composition is shown in Figure 1E, saturating at ∼20 μM total lipid. Saturation likely reflects saturation in the FRET signal rather than a saturation in fusion itself. From a standard calibration curve on GUVs prepared with known ratios of FRET donor:acceptor (Figure S1), we can calculate the number of fused liposomes (n*LUVs*) to the GUVs (see Methods for calculations), which increases from a few thousands to a few hundreds of thousands depending on liposomal concentration (Figure 1F).

Because of their large size, the GUVs contain a nearly symmetrical distribution in lipid number between their inner and outer leaflets. In contrast, due to geometric constraints, small liposomes inherently exhibit transbilayer asymmetry, with more lipids in the outer (N_out_) than the inner (N_in_). Knowing n*LUVs* and the respective liposome and GUV areas, and assuming an average GUV size of 30 μm, the calculated theoretical transbilayer asymmetry (N_out_/N_in_, see Methods for derivation) in the GUVs as a function of liposomal concentration increases from 4% up to 10% (i.e. 4-10% more lipids in the outer than the inner leaflet). Such asymmetries induce spontaneous curvature and membrane remodelling, with GUVs exhibiting outward buds; bud directionality is compatible with an excess of lipids in the outer leaflet (Figure S2). Given that the fluorescence lifetime of the FRET donor in the buds is identical to that in the main GUV membrane within experimental certainty, the lipid composition is at equilibration.

Notably, while content accumulation in the GUV lumen confirms successful fusion, a substantial fraction of GUVs simultaneously loses luminal content, suggesting leakage (Fig. 1E, inset). We have recently shown that GUVs can sustain fusion of a few tens of thousands of liposomes before perforation(19), showing that fusion and leakage are mechanistically distinct processes. We assessed GUV leakage using a three-colour confocal imaging, with green-labelled GUVs incubated with liposomes (labelled with DOPE-Atto647N, cyan) in the presence of sulforhodamine B (SRB, red), a small, membrane-impermeant fluorescent dye that served as an indicator of pore formation (44). We prepared GUVs with a composition that mimics the average plasma membrane, consisting of equimolar amounts of DOPC, DOPG, DOPE, and cholesterol(8). Without liposomes (hence no cyan signal), SRB is always retained in the outer medium (Figure 1G, top row). SRB is detected at increasing concentrations in the GUV lumen as liposome concentrations increase (Figure 1G), demonstrating progressive membrane permeabilization. Both fusion and leakage are confined to the membrane in direct contact with liposomes, as smaller vesicles occasionally present in the GUV interior are not subject to either fusion or leakage (Figure S3).

We quantify membrane permeation from the GUV degree of filling(51), where we defined values ≤ 0.1 as intact, whereas values above are defined as permeable. Figure 1H shows that permeabilization linearly increases with fusion efficiency. Irrespective of LUV concentration, GUVs exhibit a graded leakage response (in contrast with an all-or-none mechanism, where vesicles are either fully permeabilized or remain intact), consistent with transient, size-selective pores(52). Supporting this, delayed addition of a second fluorescent probe of similar size failed to enter previously permeabilized GUVs in many cases (Figure S4), showing that pores formed in GUVs are transient, eventually closing. Notably, GUVs produced with the expected composition after fusion are always intact (Figure 1H, inset and (49)), demonstrating that membrane permeabilization is not caused by changes in the lipid composition, but instead is a result of fusion-induced membrane asymmetry.

The leakage pores formed in GUVs upon liposome fusion are below the optical resolution. To estimate pore size, we fused the GUVs with increasing concentrations of liposome in the presence of a small (∼0.5 nm), medium (∼1.5 nm) or a large (∼3.6 nm) leakage probe(53), where the dimensions represent hydrodynamic radii. The extent of permeation increases with liposomal concentration for the small and medium probes, but it is always much higher for the small probe (Figure S5); i.e. as the liposomal concentration increases, the leakage pores widen. The large probe does not permeate into the GUVs, indicating that these pores are < 3.6 nm.

Liposome fusion increases GUVs tension due to number asymmetries by modifying ΔA_0_ (eq. 2), as evidenced by the suppression of membrane fluctuations. Because tension reduces the energy barrier for pore formation (eq. 1), we calculated the tension that arises due to curvature effects. This so-called spontaneous tension (σ_m_), given by *σ*_*m*_ = 2*κm*^2^, where *m* is the membrane spontaneous curvature and *k* is the bending rigidity(48) can be obtained from the diameter of the formed buds (D_bud_). For D_bud_ ranging from 0.6-2.5 μm (Figure S2 and (54)), and *m* = 2/D_bud_ (0.8 - 6 μm^-1^ for the buds here), and using *k* as 45 k_B_T for charged GUVs(55), the upper range of σ_m_ is ∼ 13 μN/m. This is well below the reported lysis tension (∼ 1 mN/m) for fluid membranes(41, 56), and the increase in tension does not explain the formation of pores in membranes exhibiting number asymmetries.

### Liposome fusion reduces membrane edge tension

Membrane stability against pore formation reflects a balance between surface tension (*σ*), which favours pore opening, and edge tension (*γ*), which resists it (Fig. 2A). In toroidal pores, where lipids bend at the rim so that headgroups line the aqueous edge while shielding hydrophobic tails, this rearrangement incurs a line energy per unit length(36, 57). Observation of leakage pores in GUVs following liposome fusion suggests that fusion transiently lowers *γ*. To directly measure the effects of fusion on γ, we incubated GUVs with 10 μM liposomes (lipid concentration) - a concentration that does not yet spontaneously permeabilize these GUVs - and measure γ from the dynamics of pore formation upon GUV electroporation(41, 58). Pores formed in control GUVs close within a few tens of milliseconds (Figure 2B, upper row and Video S1), in agreement with prior reports(43, 44). In contrast, while some GUVs incubated with liposomes exhibit qualitatively similar electroporation dynamics (Figure 2B, ii), ∼14% (4 out of 28) of the investigated GUVs collapse (Figure 2B, iii and Video S2), converting the (quasi-)flat membranes into lipid tubes, a process identical to the bursting process observed in conditions of extremely low γ(59). In rare cases, liposome fusion results in the spontaneous partial or complete collapse of GUVs; i.e. without the application of an external force (Figure 2C). Conversion of membranes into tubes indicates an increase in spontaneous curvature(60), compatible with changes in σ_m_.

**Figure 2.**
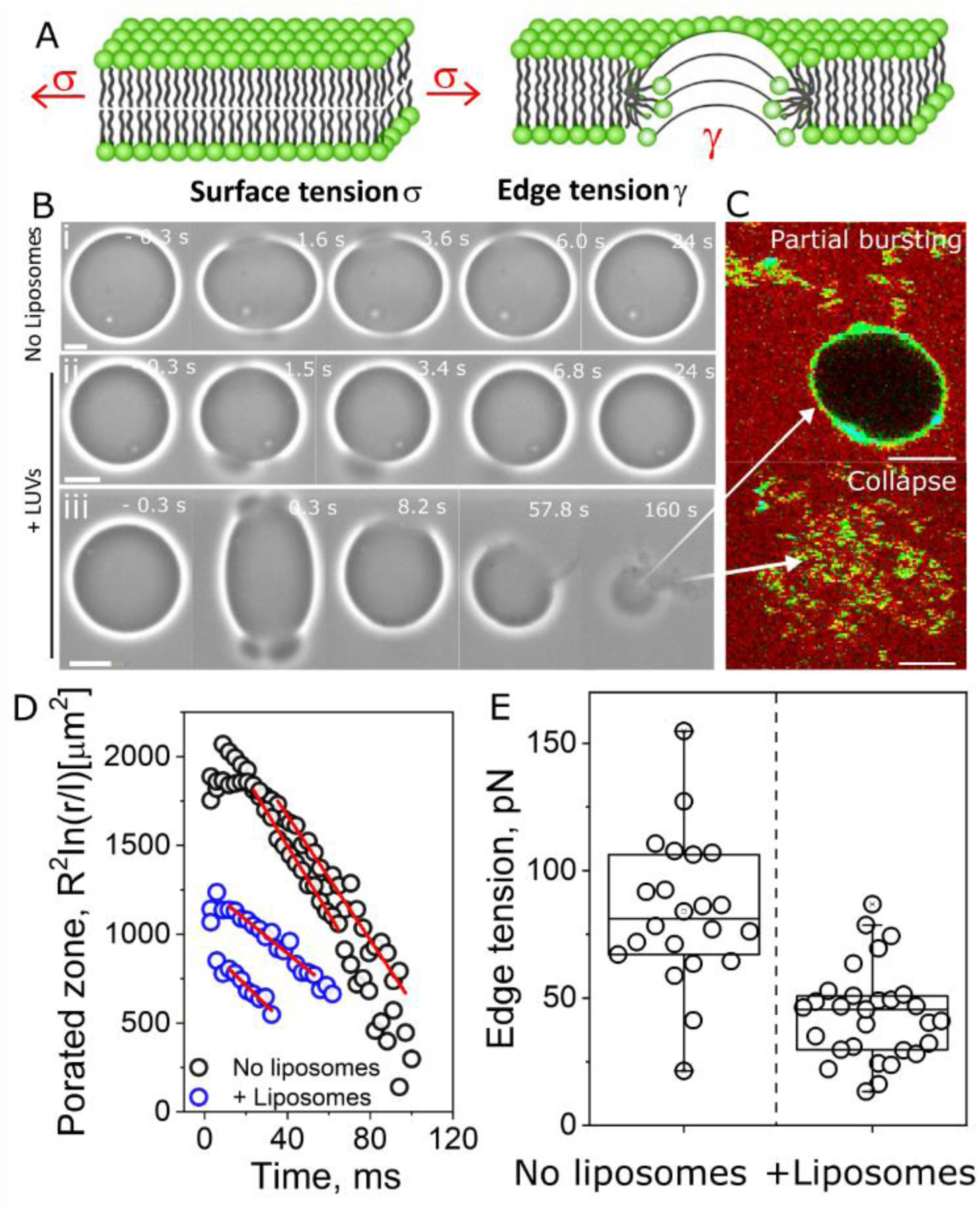
Edge tension measurements on GUVs. A, sketch of an intact bilayer upon the application of an external tension, increasing the bilayer surface tension (σ), eventually leading to the opening of a membrane pore. Due to the edge tension (γ), the formed pore spontaneously closes. B, representative DOPC:DOPG:DOPE:Chol (equimolar fractions) GUVs upon the application of a (3-4 kV/cm pulse strength and 15-200 μs pulse duration) electric field for a control GUV (no liposomes, upper row), and for a GUV that had been incubated with 10 μM liposomes (lipid concentration). The times correspond to the onset of pulse application. Field direction is indicated. C, representative GUVs (DOPC:_B_PS, 50:50 molar fraction – labelled with 0.5 mol% Bodipy C_16_) that spontaneously open pores upon liposome incubation in the presence of 0.02 mg/ml KU530. The upper and bottom images depict a GUV that undergoes partial bursting, and a GUV that fully collapses. All scale bars: 10 μm. C, temporal evolution of pore sizes formed in DOPC:DOPG:DOPE:Chol GUVs upon electroporation. Experimental data (open circles) and fit to equation 11 (red lines) are shown. E, measured edge tension from pore closure dynamics. Each point represents a measurement on an individual GUV.

For control GUVs (no liposomes), the measured γ was 84±29 pN, higher than values for neutral (42, 49) or negative GUVs(59) rich in PC lipids, likely due to the relatively high cholesterol and DOPE content. For the GUVs incubated with liposomes that survived electroporation, γ is significantly reduced to 44±18 pN. This shows that the asymmetry induced from liposome fusion lowers edge tension. Upon pore formation, closure occurs only if γ is ≥ 5-10 pN(59); lower or even negative (61) values result in membrane collapse. Because these measurements are limited to GUVs that survived electroporation, they likely underestimate the full effects of fusion, as a significant fraction of GUVs collapse. Overall, the results show that transbilayer asymmetries that originate from liposome fusion compromises membrane stability by reducing edge tension, consistent with the spontaneous perforation of GUVs and, in extreme cases, complete vesicle collapse.

### Membrane fusion generates transient pores of defined size in living cells

To test the generality of our observations, we incubated the fusogenic liposomes with living cells. HEK cells were incubated with LUVs (500 μM lipid concentration) for 30 minutes in the presence of two probes of different molecular weights: SRB (∼ 0.5 nm, red) and Dextran 10 kDa (∼3.6 nm, green). The elevated LUV concentration compared to GUVs reflects the combination of high cell density and a greater excess membrane area per cell (54). Following the GUV results, we hypothesize that the formation of small pores would enable the entry of SRB but not Dextran 10 kDa in the cells (Figure 3A). Without liposomes, both probes are retained in the outer medium (Video S3). Figure 3B shows a cell that becomes permeable to SRB upon liposome addition but not to Dextran 10 kDa, surrounded by cells intact to both probes. Intensity line profiles (Figure 3C) show that the intracellular signal is well above background for the permeable cell due to SRB affinity for cytosolic structures, as previously observed(62). Figure 3D shows the degree of filling for several individual cells after 30 minutes incubation. For SRB, cell permeabilization is an all-or-none process; once permeable, entry is always complete, showing that the pores remain open at least until SRB full permeation. In contrast, most of these cells remain intact to Dextran 10 kDa, and those that do become permeable, only exhibit graded permeabilization, suggesting the formation of initially larger pores that eventually shrink.

**Figure 3.**
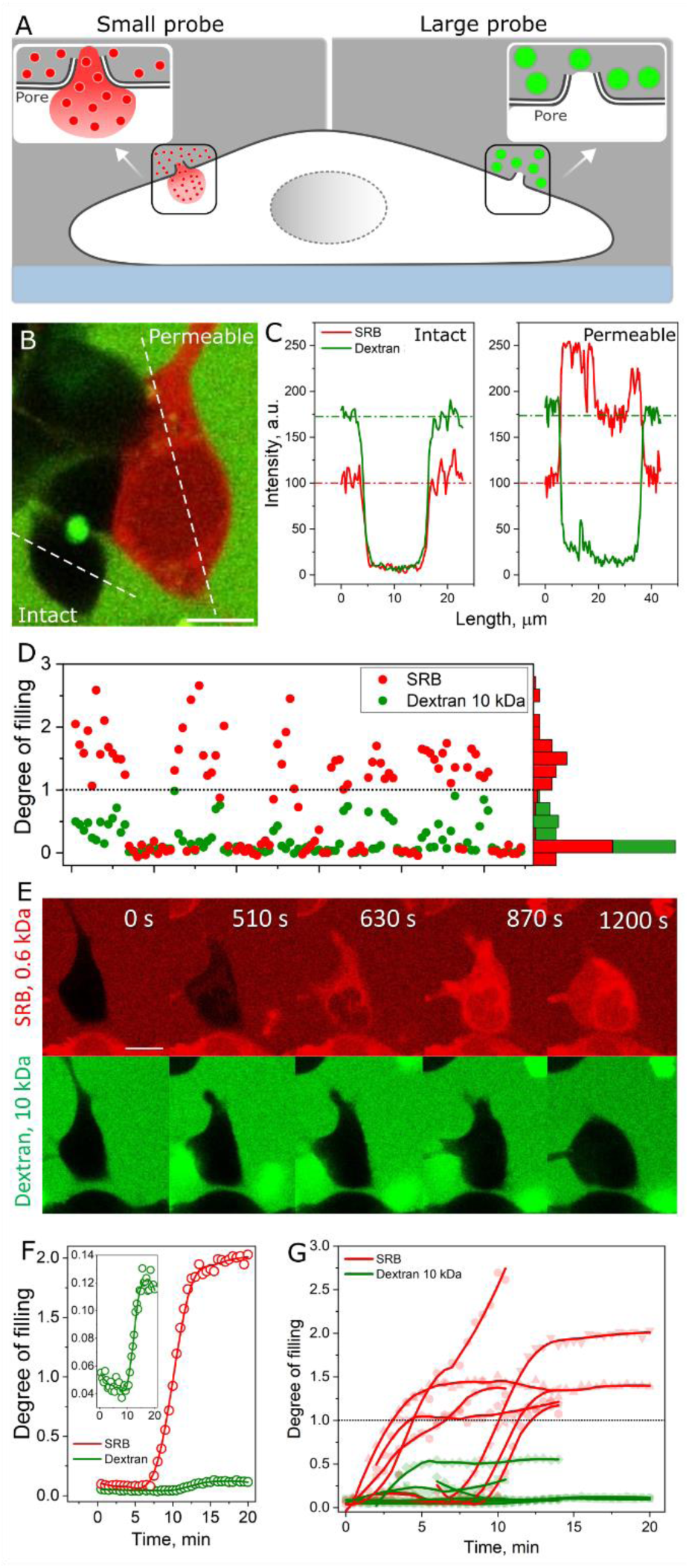
Liposome fusion opens transient leakage pores of defined sizes in living cells. Sketch of pore formation and leakage of a small (red) and large (green) probe into the cell. B, representative image of a HEK cell that is permeable to the small probe SRB (5 μM in the outer medium) but not permeable to the larger probe Dextran 10 kDa (0.1 mg/ml in the outer medium). Note that the cell is surrounded by non-permeable cells (black interior). C, intensity line profiles from the lines in B for SRB (red) and Dextran 10 kDa (green) in an intact cell (left) and in a cell permeable to SRB only (right). D, degree of filling on many individual HEK cells for SRB and Dextran 10 kDa. Each point represents a measurement on an individual cell. The plot also shows the histogram distribution. Hatched line: degree of filling = 1 (full permeation). Higher values for SRB are a result of the probe affinity for intracellular structures. In B-E, the cells had been incubated with 500 μM liposomes (lipid concentration) for 30 minutes before imaging. E, permeation kinetics of SRB and Dextran 10 kDa in a single HEK cell upon incubation with 500 μM liposomes (lipid concentration). The numbers correspond to the time from the onset of observation. F, representative permeation kinetics of both probes in a single HEK cell. Inset: zoom-in of Dextran 10 kDa. G, permeation kinetics on several (7 individual cells from 2 independent experiments) HEK cells. All scale bars: 10μm.

To gain information about pore dynamics, real-time permeabilization experiments (Figure 3E and Video S4) show the permeabilization of a single HEK cell that becomes fully permeable to SRB and allow some entry of Dextran 10 kDa (kinetics in Figure 3 F). SRB permeation develops until completeness within ∼7 minutes for this cell (6 ± 2 min for n = 7 cells from two independent experiments), whereas Dextran 10 kDa permeation is very modest (degree of filling ∼ 0.12) and short-lived (∼ 5 minutes). Permeabilization is accompanied by shape remodeling, indicating that pore formation is toxic to cells. This behaviour is highly reproducible, with pores opening stochastically (Figure 3G). A close examination of the permeabilization dynamics shows a delay of 3 ± 2 min between SRB and Dextran 10 kDa entry, suggesting that first small pores open allowing only SRB entry, then they expand into larger pores allowing entry of Dextran 10 kDa (presumably as the liposomes continue to fuse with the cells), then they shrink stopping Dextran 10 kDa flow. In fact, leakage pores open in HEK cells and eventually close(62). Considering SRB size of 0.5 nm and diffusion coefficient of 400 μm^2^/s in the outer solution(63), with full permeation within 5 minutes, and assuming that the membranes contain pores of 2 nm diameter (slightly smaller than the lower bound of Dextran 10 kDa diameter), the plasma membrane of HEK cells contains ∼ 10 pores. In summary, the results show the opening of (multiple) transient pores of variable size that eventually close, in qualitatively agreement with pores formed in GUVs.

### Pores open spontaneously in membranes with lipid number asymmetries

The experiments show that liposome fusion induces spontaneous, transient pores of defined size, driven by reduced edge tension in both model membranes and living cells. We propose that transbilayer lipid-number asymmetries sensitize bilayers to pore formation, reducing edge tension, although the mechanism of subsequent pore closure remains unclear. To probe the molecular basis of pore opening and sealing, we performed coarse-grained (CG) molecular dynamics simulations with the Martini force field. However, on the accessible sub-millisecond timescale of the simulations, even strongly asymmetric bilayers remain highly stable because the bilayer becomes kinetically trapped in a metastable state (64). As a result, spontaneous pore formation is rarely observed for typical phospholipids with c16 – c18 tails. In contrast, lipids with shorter hydrocarbon tails reduce membrane thickness and cohesion, lowering the line tension at pore edges. This enables stress generated by leaflet asymmetry to be released through spontaneous pore nucleation, amenable to CG simulation.

To this end, we simulated flat membrane patches composed of the short-tail lipid 1,2-ditricosanoyl-sn-glycero-3-phosphocholine (DTPC) with progressively increasing lipid-number asymmetry across the bilayer. Starting from symmetric membranes containing 2500 lipids per leaflet, we gradually removed lipids from the inner leaflet, mimicking the excess lipids in the outer leaflet observed in the liposome and cell-fusion experiments. Simulations were performed under constant-area conditions to prevent membrane buckling. Membranes with no or low (< 14%) asymmetry remained stable throughout the simulations (Figure 4A). At 16% asymmetry or higher, pores spontaneously (Video S5) and stochastically (Figure S6) formed in these membranes. As transbilayer asymmetry increased, the spontaneous pores became larger and formed earlier (Figure 4B). These results demonstrate that transbilayer lipid number asymmetries alone are sufficient to induce spontaneous pore formation in membranes, and support the widening of pores formed in cells as more liposomes fuse (thus delivering greater number asymmetries).

**Figure 4.**
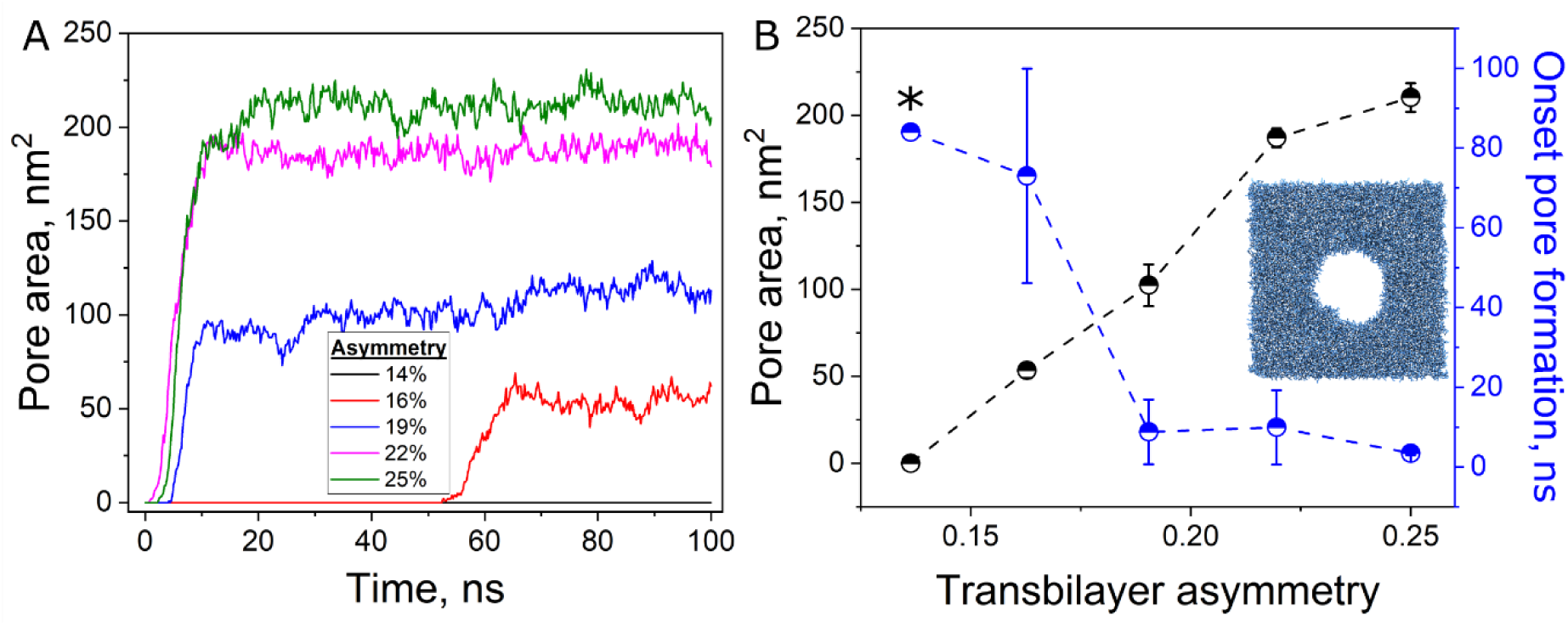
Spontaneous pore formation in membranes containing transbilayer lipid number asymmetry. Molecular dynamic simulation of a 40×40 nm membrane patch made of DTPC lipids containing 2500 total lipids in the upper leaflet and increasingly lower lipid number in the lower leaflet. A, pore area for membranes with various transmembrane asymmetries. B, area (black) and onset of pore formation (blue) for pores formed in bilayer patches as a function of transbilayer asymmetries. The asterisk at 14% asymmetry refers to only one of three simulations leading to pore opening, whereas a pore did not open in the other two simulations. The data shows the mean and s.d. for three runs. Inset: a pore in a bilayer patch containing 2500:1900 lipids (upper:lower leaflet).

### Number asymmetry is relieved via flip flop through the pore, catalysing pore closure

Pores formed due to transbilayer lipid asymmetries spontaneously close in both biomimetic and living cell membranes, despite markedly different surface tensions. In unsupported GUVs, resting tension is low (∼10⁻⁹–10⁻⁶ N/m)(65, 66), whereas in living cells, it is significantly higher (10^-3^-10^-2^ pN/m(67)). This difference likely accounts for the substantially longer pore lifetimes observed in cells(68, 69) compared to GUVs(43, 44). Membrane support not only increases tension, but it also favours the persistence of open pores(70). To probe pore dynamics under these contrasting conditions, we performed CG MD simulations of membrane patches using two setups: a constant-surface-tension ensemble (laterally compressible membrane, mimicking GUVs), and a constant-area condition (laterally constrained membrane), in which the bilayer area is constrained in the lateral dimensions, thus permitting a build up in tension, to mimic mechanically supported cell membranes.

We used membranes made of equimolar amounts of DOPC:DOPE:DOPG:Chol. Because pores do not form spontaneously in these membranes, even on the timescale of 100 microseconds (Figure S7), we artificially created a single pore of 6 nm diameter by gradually applying a cylindrical restraint to the membrane centre, displacing lipid tails and expelling them from the pore region. The hydrophilic heads are not restrained in order to allow lipid flip flop through the pore. The cylindrical constraints are applied for a fixed period of 100 ns. In the laterally compressible setup (the membrane is allowed to relax), the opened pores have a constant area ∼ 27 nm^2^ and spontaneously close as soon as the constraints are relieved, irrespective of transbilayer asymmetry (Figure 5 A-C and Videos S6 and S7). In contrast, while pores formed in laterally constrained membranes at low asymmetry (up to 15%) spontaneously close (Video S8), higher asymmetries lead to stably open pores for as long as the simulation runs (Figure 5B, S8A and S8B and Video S9). Similarly to spontaneously opening pores, externally generated pores exhibit increased size and lifetime as asymmetry increases.

**Figure 5.**
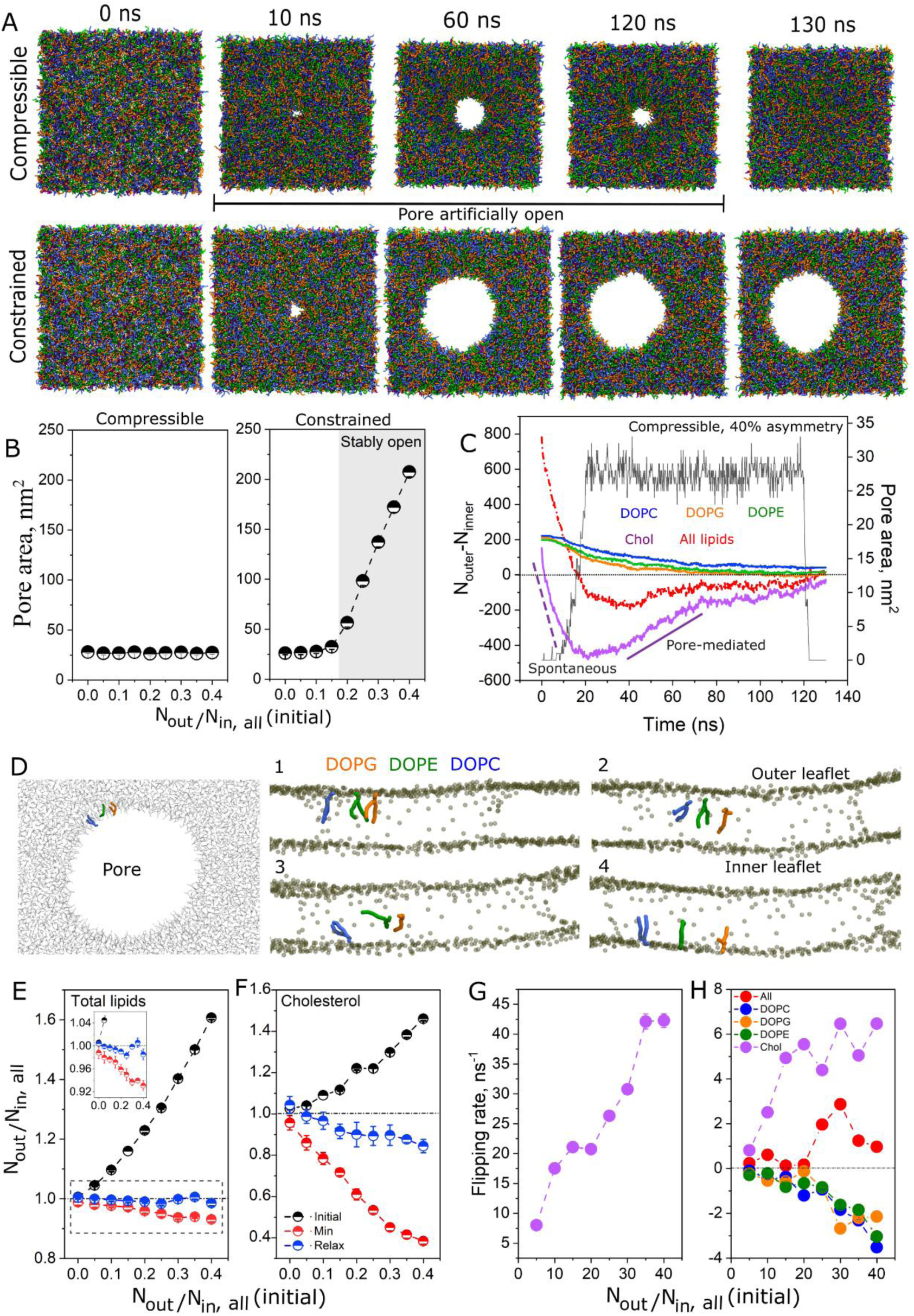
Pore formation alleviates transbilayer asymmetries. A, representative snapshots of a MD simulation of a DOPC:DOPE:DOPG:Chol (equimolar fractions) membrane patch exhibiting 40% number asymmetries at different timepoints for both the compressible and the laterally constrained membrane. The pore is kept artificially open from 10 ns to 120 ns. B, equilibrium pore area as a function of initially imposed number asymmetry for compressible and constrained membranes. The grey bar indicates stable pore opening. C, lipid difference in lipid number in the outer and inner leaflets for a membrane exhibiting 30% number asymmetry in laterally constrained membranes. The region where the pore is open is also shown (black data). Purple lines show the linear fits of the fast flip flop regime for cholesterol for spontaneous flip flop before pore opening (hashed line) and cholesterol’s redistribution upon pore opening (solid line). D, representative snapshots of lipid flip flop from a top view (left image) and lateral view of the pore (right images) upon pore formation. One representative molecule of each phospholipid type is highlighted. For the remainder of the membrane, only the phosphate beads are shown, highlighting those lipids lined at the pore. Flip flop time is completed within ∼ 20 ns. E-F, N_out_/N_in_ as a function of initial (pre-defined) asymmetries for all four lipids (Figure E) and for cholesterol only (Figure G) before lipid equilibration (initial, black data), upon full equilibration and before pore opening (min, red data) and after relaxation upon pore opening-closure (relax, blue data). Each data point represents the mean and s.d. from three independent simulations. The inset in E shows a zoom-in at lower asymmetries. G, measured rate of spontaneous flip flop for cholesterol before pore opening. Measured flip flop rate for all lipids as a function of transbilayer asymmetries.

By plotting the temporal evolution of the difference in lipid number in the outer (N_outer_) and inner leaflet (N_inner_) for all four lipid species, it becomes evident that even before pore opening (t < 20 ns), the initial transmembrane asymmetry progressively vanishes—and even overshoots to values below 1, as lipids accumulate in the lower leaflet. Note that this overshooting is driven by rapid flip flop and accumulation of cholesterol in the lower leaflet (Figure 5C, Video S10), balancing area mismatches; since the area of cholesterol is smaller than the area of the phospholipids, it is required in larger numbers to balance such mismatches. In fact, cholesterol flip flop is fast and occurs within a coupe of ns in intact membranes (Figure S9). The extent of overshooting depends on the initial magnitude of asymmetry, featuring greater extents of cholesterol flip flop at larger asymmetries, as shown for compressible (Figure 5E-F) and constrained (Figure S8C-E) membranes. Note that the balancing in leaflet areas comes at the expense of generating compositionally asymmetric membranes, with cholesterol being highly depleted from the upper leaflet. In contrast to cholesterol, none of the phospholipids exhibit flip flop in intact membranes (Figure S10).

Interestingly, pore opening allows phospholipids to flip through the pore (Figure 5D and Video S11). Initially, phospholipid headgroups align along the pore rim, visible along the phospholipid headgroups shown in the snapshots, after which they undergo a transverse reorientation while their hydrophobic acyl chains remain within the bilayer core, ultimately enabling their relocation to the inner leaflet within ∼ 20 ns. This is mechanistically identical to the credit card model of lipid scramblase-mediated lipid flip flop(71). Note a clear directionality, from the initially overpopulated outer leaflet to the inner leaflet. Figures S11 and S12 show the dynamics of asymmetry relieve across the whole asymmetry range for laterally compressible and laterally constrained membranes, respectively. In Figure 5E, black data represent the asymmetry ratio N_out_/N_in_ at the beginning of the simulations (reflecting the imposed asymmetry) prior to both spontaneous lipid redistribution and pore formation, with an excess of lipids in the upper leaflet. Before pore opening and at 25 mol% cholesterol, lipid number asymmetries are fully relaxed via cholesterol flip-flop for initial N_out_/N_in_ differences <5% (red data). Larger initial asymmetries leave increasingly larger residual imbalances. This buffering effect, although significant, is weaker than that of 10 mol% CerC_6_(72), likely due to cholesterol’s deeper penetration in the membrane and its lower effective curvature relief. Because this occurs before pore opening, results are similar in laterally compressible and laterally constrained systems (Figure S8).

Upon pore opening, transmembrane asymmetries largely disappear as the lipids flip flop through the pore (cholesterol flip flop occurs away from the pore) along their concentration gradients. This leads to nearly symmetric membranes despite initially large asymmetries (blue data in Figure 5E). This phospholipid redistribution restores cholesterol’s initial distribution (Fig. 5E, Video S12) because, in intact membranes, cholesterol flips toward the stretched leaflet to buffer asymmetries, whereas pore-mediated phospholipid flip flop reverses this effect. These dynamics occur in both laterally compressible and constrained systems (Fig. S8), indicating that cholesterol’s leaflet preference is driven by differential stress rather than chemical affinity in our system. Notably, in laterally compressible membranes, residual cholesterol enrichment persists for initial asymmetries >10% (blue data, Fig. 5H) because pores close before full equilibration. In contrast, pores that remain open (i.e. in constrained membranes) allow complete relaxation, with cholesterol becoming evenly distributed (Figs. S8E, S12). We conclude that pore opening promotes lipid flip flop, effectively relaxing number asymmetries in a manner dependent on initial imbalance and pore lifetime, and restoring cholesterol’s original distribution.

Because pore opening is driven by transbilayer asymmetry (Eq.2), lipid flip flop through the pore rapidly reduces these asymmetries to values below the threshold required for pore formation. As the asymmetry decreases, the energetic cost of maintaining the pore rises as ΔA/A₀ lowers - and the edge tension likely increases as well - promoting pore closure. Thus, pore opening provides a rapid physical pathway for equilibrating leaflet asymmetries, dissipating the very tension that drives pore formation—making pore opening intrinsically self-limiting.

### Flip flop rates through the pore

A toroidal membrane pore connects both leaflets, enabling lipids to translocate between them, a process analogous to lipid scrambling. Like scramblases (46), this pore-mediated flip flop is non specific and energy-independent. For scramblases, scrambling rates are in the order of 10^3^ s^-1^(73) to 10^6^ s^-1^ (74), in contrast to their slower counterparts flippases and floppases, which require ATP-dependent conformational changes, are often lipid-specific, and operate at much slower rates (10-100 s⁻¹)(75, 76).

Lipid distribution in asymmetric membranes is governed by spontaneous(72, 77) cholesterol flip flop before pore opening (or by other molecules capable of flipping (72, 77)), and by the flip flop of all lipid types once a pore forms, restoring cholesterol to its original distribution. Directional lipid transport from the outer to inner leaflet was quantified by fitting the initial linear regime of lipid number differences over time (Figure 5C). Before pore formation, flip flop occurs exclusively via spontaneous cholesterol translocations, with rates increasing with initial transbilayer asymmetry and reaching remarkable values on the order of tens of flip flops ns⁻¹ (Fig. 5G). After pore opening, all lipids can translocate, with flip flop rates about an order of magnitude lower than cholesterol but still in the ns⁻¹ range (Fig. 5H); 4–8 orders of magnitude faster than known protein-mediated mechanisms (10³–10⁶ s⁻¹). Rates are non-specific, with all lipids showing similar kinetics (the sign indicates flip flop directionality), scaling with initial asymmetry.

### Lipid composition in intact membranes and in the pore region

Once a pore opens, the membrane is no longer laterally or geometrically uniform. The pore edge constitutes a localized, highly curved defect that drives curvature-dependent lipid sorting(78). A lipid pore is surrounded by a toroidal rim characterized by extreme curvature and packing frustration, causing lipids to redistribute according to their molecular shape(79). Lipids with curvature compatible with the pore rim are enriched, whereas inverted-cone lipids with negative spontaneous curvature are depleted into flatter membrane regions; cylindrical lipids are largely curvature-neutral and redistribute weakly, mainly driven by the affinity of the other lipids. This sorting is expected even in membranes that are otherwise compositionally and numerically symmetric.

We performed MD simulations to spatially and temporally resolve the distribution of lipids upon pore formation in DOPC:DOPE:DOPG:Chol at 0 and 40% number asymmetries to reveal the chemical composition of intact membrane regions and pore rims in both symmetric and number-asymmetric membranes. Given its cylindrical shape, DOPC is not expected to be particularly enriched in the curved pore rim. Due to its charged headgroup, DOPG is strongly hydrated, effectively increasing its headgroup area(80). Together with electrostatic repulsion(36), this is expected to promote enrichment of DOPG in pore regions, explaining the reduced edge tension in the DOPG membranes(59, 81). In contrast, the inverted-cone shape lipids DOPE and cholesterol are expected to be excluded from the pore (i.e. accumulated in regions of intact membrane).

Figure 6A shows representative snapshots of a DOPC:DOPE:DOPG:Chol bilayer exhibiting 40% number asymmetry and a toroidal pore. Since all four lipids are present at 25 mol%, a local fraction of 0.25 indicates a uniform distribution, while deviations from this value reflect lipid enrichment or depletion. Figures 6B shows the dynamic fractions of each of the lipids at the pore rim (within 5 nm from the pore edges) during pore-mediated lipid redistribution in a membrane exhibiting 40% asymmetry, and Figure 6C indicates the average lipid distribution. The lipid distribution in intact regions of the membrane is shown in Figure S13. The results show lipid-specific enrichment/depletion patterns in membranes containing a pore. Cholesterol is enriched in intact regions (Fig. S13A,B), driven by its strong depletion from the pore rim (Figure 6B,C)—consistent with its inversed-cone shape and associated curvature preference. In contrast, DOPE exhibits virtually no pronounced partitioning preference despite its inversed-cone shape. DOPC and DOPG are depleted from intact regions due to their accumulation at the pore rim. We interpret this enrichment as a secondary effect of cholesterol’s strong exclusion from the pore region, effectively “pushing” these lipids toward the pore rim.

**Figure 6.**
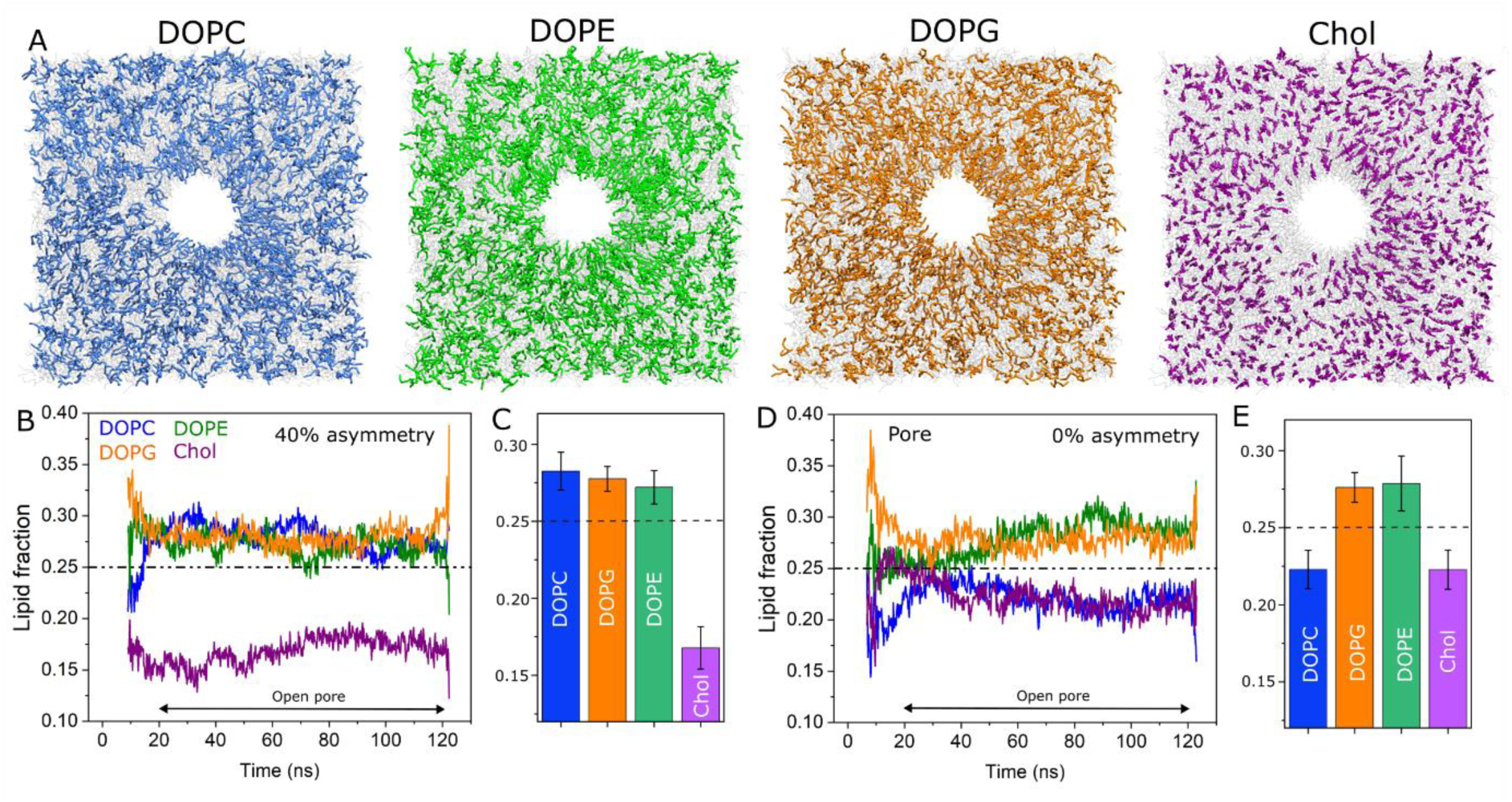
Pore-mediated lipid sorting is enhanced by asymmetry. A, representative snapshots of a pore in DOPC:DOPE:DOPG:Chol (equimolar fractions) membrane exhibiting 40% number asymmetry. The snapshots are taken before pore closure. B,D: dynamic fraction of lipids as a function of time within 5 nm the pore rim at 40% (in B), and 0% asymmetry (in D). The time window where the pore is open (t = 20–120 ns) is indicated. C and E, average fraction of lipids around the pore for membranes exhibiting 40 % (in C), and 0% asymmetry (in E). The horizontal hatched lines at 0.25 represents lipids’ expected distribution in the absence of compositional preferences.

In a symmetric membrane (i.e. 0% asymmetry, Figs. 6 D,E and S13C,D), lipid sorting is much weaker. DOPE and DOPG are slightly enriched around the pore, whereas DOPC and cholesterol are mildly depleted. The local accumulation of lipids that lower edge tension (i.e. PG(59) and PE(41)), along with the depletion of those that raise it (i.e. cholesterol (42, 49)), may help explain the reduced stability of membranes enriched in charged lipids or DOPE(41).

Notably, our results show that, in addition to transbilayer number asymmetry that builds during fusion, which results in a strong compositional asymmetry driven by enrichment of cholesterol in underpopulated leaflet, pore formation itself generates local compositional heterogeneities, all of which are known to modulate membrane structure and stability. Pores act as transient sorting domains that selectively enrich or deplete specific lipids, with partitioning dependent on lipid type and magnitude of leaflet asymmetry: for example, the cholesterol fraction near the pore decreases by ∼40% in highly asymmetric membranes but by only ∼12% in symmetric ones. Lipid redistribution through the pore relaxes leaflet number and compositional asymmetries, removing the driving force for pore formation, thus enabling resealing. Because membrane composition directly determines the material properties that regulate pore dynamics and stability, its regulation by leaflet number asymmetry, flip flop–capable molecules, and pore opening renders these asymmetry-induced effects functionally significant.

## DISCUSSION

The substantial metabolic investment required to maintain the plasma membrane far from equilibrium underscores the importance of lipid asymmetry between the two leaflets. Cells must suppress pore formation to preserve lipid-number and compositional asymmetry. Physically, membrane growth must remain sustainable: asymmetric lipid insertion—via synthesis or fusion—creates leaflet area mismatches, elevates surface tension, and reduces edge tension, potentially resulting in membrane rupture and formation of hydrophilic pores. Lipid accumulation within a single leaflet can occur on minutes timescales in various biological processes, such as bacterial growth (46) and growth of synthetic cells(19)—requiring equally rapid relaxation mechanisms to prevent membrane failure. Flip flop–capable molecules provide a first layer of protection by buffering moderate asymmetries through spontaneous flip flop(16). Beyond this, pores form: if they remain below the critical radius r*, they are transient and the lifetime is regulated by edge and surface tension (lower in GUVs and higher in cells); larger pores (r > r*) cause catastrophic rupture. In this view, pores function as lipid scramblases—flipping molecules prevent pore nucleation at moderate asymmetries, and once pores appear, they remain below r* long enough for lipid asymmetries to relax via flip flop through the pore, after which edge tension drives closure. In this picture, edge tension is likely not a static but a dynamic parameter that depends on the magnitude of asymmetry, and increases as lipids flip through the pore.

Flipping molecules can only compensate lipid number/area asymmetries up to a finite limit, determined by their concentration, the magnitude of the asymmetry and the location within the bilayer(72). This constraint helps explain why cholesterol is unusually abundant in the PM (∼30–40 mol(5, 6, 8, 9)). However, cholesterol also alters bulk membrane properties—raising viscosity(49) and reducing flexibility, especially in membranes rich in long, saturated lipids (82). Thus, adjusting asymmetry via cholesterol has dual effects on both asymmetry and bulk mechanics. Consequently, cholesterol cannot reach similar levels in all membranes; for example, ER membranes contain only ∼5 -15 mol % cholesterol(83), making spontaneous cholesterol flip flop insufficient to buffer large asymmetries during rapid organelle growth. A similar limitation applies to synthetic cells: achieving the twofold area increase required for symmetric division would generate extremely large asymmetries, rendering the membrane mechanically unstable(19).

For a lateral tension applied on asymmetric membranes that originates from curvature effects, the corresponding curvature-induced tension (σ_m_) can be estimated from the bud diameter D_Bud_ as σ_m_ = 2*k*m^2^, where *κ* is the bending rigidity (*κ* ≈ 45 *k*_B_*T* (55)) and *m* is the spontaneous curvature *m* = 2/*D*_bud_(48). For the observed buds here, this yields *m* ≈ 0.8-6 μm^-1^ and corresponding σ_m_ ≈ 10-15 μN/m, well below the reported lysis tension of fluid lipid membranes (∼ 5-10 mN/m(56)). Thus, curvature-induced tension alone cannot account for the observed GUV destabilization. From eq. 1, membrane stability is given by the balance between lateral and edge tension, and as shown by the results, a decrease in edge tension reduces membrane stability.

Within the area-difference elasticity (ADE) framework, the relaxed area difference is given by ΔA_0_/A = 2h*m* (84, 85), where h is the monolayer thickness (here, 2 nm). For the curvatures observed, this yields an upper bound ΔA_0_/A ≈ 2.5%. For DOPC:DOPG membranes at low ionic strengths and γ ≈ 5-15 pN(59), r* calculated from eq. 3 is ≈ 0.5-3 nm. In contrast, for the GUVs studied here rich in chol and DOPE, γ ∼ 85 pN and r* ≈ 30 nm prior to LUV fusion, decreasing to ≈ 10 nm after fusion (for γ ≈ 45 pN). The lipid number asymmetry inferred from r* using Eq. 3 is Δ*A*/*A* ≈ 5%, slightly larger than the ADE-based estimate (≈ 2.5%) and comparable to the experimentally measured value (≈ 4-10%). These results demonstrate a direct coupling between leaflet asymmetries area and membrane stability, as reflected in the reduction of *r*\*.

If edge tension decreases sufficiently, the pore nucleation barrier Δ*G*^∗^becomes comparable to thermal energy, while *r*^∗^shrinks to molecular length scales, resulting in the spontaneous collapse of membranes. This is consistent with permeability measurements at 10 µM LUVs, where sub-nanometer probes (KU530) permeate the membrane, whereas larger probes (10 kDa) do not, indicating pores smaller than 2.5 nm, in agreement with an estimated r* ≈ 4.5 nm under these conditions, as well as spontaneous GUV bursting at higher LUV concentrations (Fig. 2C). We conclude that sufficiently large lipid number (or area) asymmetries greatly reduce edge tension, enabling spontaneous pore nucleation and mechanical instability even under vanishingly small lateral tension.

Beyond regulating lipid number asymmetries, membrane pores can be viewed as curvature-sorting regions imposed by molecular shape(36, 57). Lipids with negative spontaneous curvature are expected to be depleted from the pore edge, whereas cylindrical lipids are curvature-neutral and should exhibit a partition that is dominated by the curvature-active molecules (e.g. DOPE and chol). Curvature–composition coupling further implies that even at identical bulk fractions, lipids with stronger curvature preference are enriched at the pore rim, outcompeting weaker ones. This is clearly shown for chol’s strong depletion from the pore region, “pushing” the other lipids to the pore. Treating the pore as a short-lived conduit, lipid flip-flop becomes diffusion-limited(34) and strongly driven by the leaflet area imbalance(20, 21), similarly to a lipid channel (i.e. scramblase).

At moderate asymmetries, pores do not open spontaneously and chol spontaneous flip flop efficiently relaxes number asymmetries up to ΔA/A ∼ 5%(72). Larger initial asymmetries are not fully buffered, leaving a residual number imbalance. Transient opening of a 6 nm pore for ∼ 100 ns in a 40 nm^2^ membrane patch fully dissipates asymmetries as large as 40%, corresponding to effective lipid transfer rates of 10⁹–10¹⁰ s⁻¹ and equilibration within ∼60 ns. Scaling this to a single 5-nm pore (r ≈ r*) in a 20-µm GUV yields an estimated relaxation time of ∼ 2 s for the experimentally observed 5% number asymmetry. Thus, the transient opening of a single nanometer-scale pore is sufficient to relax realistic lipid number (and compositional) asymmetries in a cell-sized compartment in a few seconds, compatible with partial rather than complete probe leakage (i.e. degree of filling < 1), restoring initial lipid distribution and effectively recapitulating scramblase function.

### Conclusion

Lipid number and area imbalances are ubiquitous in the cell. Their destabilizing effects increase membrane tension, generate curvature and remodelling, and reduce edge tension, highlighting leaflet number imbalance as a central determinant of membrane mechanics. Yet, cells are largely resistant to pore formation through dedicated protein machineries, such as lipid scramblases, rapidly alleviating such imbalances. Our results show that small pores, rather than being purely disruptive, function analogously to scramblases; they act as transient and non-specific lipid channels, alleviating both number and compositions asymmetries and catalysing their own closure. Our findings reveal that lipid number asymmetry and transient pores, fine tuned by the presence of flipping molecules, act as intrinsic regulators of membrane mechanics and morphology, providing a universal, physical mechanism for membranes to remodel, equilibrate, and preserve bilayer integrity in the absence of complex protein machineries. We hypothesize that primitive cells may have relied on a dual protective strategy combining simple flipping molecules and transient pore formation to prevent collapse under conditions of area imbalances, such as asymmetric lipid insertion and growth.

## Materials and methods

All chemicals and materials were used as obtained and without further purification. The phospholipids 1,2-dioleoyl-sn-glycero-3-phosphocholine (DOPC), 1,2-dioleoyl-snglycero-3-phospho-(1′-rac-glycerol) (sodium salt) (DOPG), L-α-phosphatidylserine (Brain, Porcine) (sodium salt) (brain PS, _B_PS), 1,2-dioleoylsn-glycero-3-phosphoethanolamine (DOPE), l-α-phosphatidylserine (Brain, Porcine) (sodium salt) (PS), 1,2-dioleoyl-3-trimethylammoniumpropane (DOTAP), and cholesterol (plant derived), as well as the fluorescent dye 1,2-dipalmitoyl-sn-glycero-3-phosphoethanolamine-N-(lissamine rhodamine B sulfonyl) (ammonium salt) (DPPE-Rh) were purchased from Avanti Polar Lipids (Alabaster, AL). DOPE-Atto647N was purchased from AttoTech (Siegen, Germany). Lipid solutions were prepared in chloroform and stored at –20 °C until use. Glucose, sucrose, Dimethyl sulfoxide (DMSO) and the fluorescent probes Bodipy C_16_ (BODIPY FL C1_6;_ 4,4-Difluoro-5,7-Dimethyl-4-Bora-3a,4a-Diaza-s-Indacene-3-Hexadecanoic Acid), sulforhodamine B (SRB), Dextran-FITC 3 kDa, Dextran-FITC 10 kDa, albumin–fluorescein isothiocyanate conjugate (FITC-albumin) were purchased from Sigma-Aldrich (St. Louis,MO, USA). Bovine serum albumin (BSA) and Poly(vinyl alcohol), PVA 87–90% hydrolyzed were also obtained from Sigma-Aldrich. The fluorescent probe KU530 NHS-Ester was purchased from KU dyes (Copenhagen, Denmark).

### Preparation and incubation of giant unilamellar vesicles and small liposomes

Giant unilamellar vesicles were prepared using the method of swelling of hybrid films of lipid and polymer(44), using nd Poly(vinylalcohol), PVA as a polymer(86). In short, ∼ 100 μL of a 2% (w/v) PVA solution in water was spread on two glass coverslips and the water was left to evaporate by placing the coverslip on a hot plate at ≈ 60 °C for approximately 10 min to form a PVA film. Next, a ≈ 10 μL of a 2 mM lipid solution containing the desired lipid mixture was spread on the PVA film. Chloroform was evaporated under a stream of Argon, and then the coverslips were sandwiched using a Teflon spacer forming a ≈ 1.8 mL chamber sealed with the help of clips. The chamber was filled with a 200 mM sucrose solution (unless stated otherwise) for ≈ 30 min to allow GUV swelling at room temperature (R.T. = 18 ± 1 °C). For lipid mixtures containing fluorescent lipids, hydration was carried out in the dark. After swelling, the GUV solution was harvested by gently pipetting the GUVs, which were used within 1 day. For imaging, the GUVs were diluted in an isotonic glucose solution to help sediment the vesicles to the bottom of the imaging chamber. If not mentioned otherwise, the GUVs were labelled with 0.5 mol% fluorescent labels.

Liposomes made of DOTAP:DOPE (50:50, molar ratio) were prepared using the hydration-sonication method(87). For the FRET experiments, the liposomes were labelled with 2 mol% DPPE-Rh, whereas for multicolour confocal experiments, the liposomes were labelled with 0.5 mol% DOPE-Atto647N. The appropriate lipid mixture in chloroform was added to the bottom of a glass chamber and evaporated under a stream of Ar and further evaporated under vacuum for 1–2 h to remove any trace of the solvent. Afterwards, the lipid films were hydrated with a 200 mM sucrose solution and vigorously vortexed until full lipid film detachment forming multilamellar vesicles (MVLs).

For the encapsulation of water-soluble probe Dextran 10 kDa-FITC, a solution containing 0.1 mg.ml^−1^ of the probe was using for hydration of 2 mM lipids and diluted ≈ 100 times, minimizing background signal. The MVLs were sonicated using a bath sonication for approximately 20 min and used within 2–3 days. GUV incubation with the liposomes was done by diluting the liposomes in isosmolar sucrose to a 100 μM lipid concentration, and an equivalent amount of the prediluted liposomes solution was mixed with 50 μL GUVs in glucose at the desired final liposomes concentration for a final 100 μL solution. GUVs and liposomes were incubated for 10–15 min in an Eppendorf tube, after which the solution was ready for imaging.

The edge tension measurements were performed as in (59, 81). In short, GUVs produced by electroformation in 200 mM sucrose solution were dissolved in isosmolar glucose solution containing either 0.1 mM EDTA or 0.5 mM NaCl, or 0.2 mM CaCl_2_. The salts were added only to the outer solution to induce mild square-oblate GUV deformation (88). Electroporation was performed using a DC electric field through a Multiporator (Eppendorf) with a 150-200 V (3-4 kV/cm) pulse strength and 150-200 μs pulse duration, connected to an electrofusion chamber (Eppendorf, Germany) containing two parallel cylindrical electrodes (92 mm radius) spaced by 500 µm. The images were acquired using a 40X (0.75 NA) air objective and imaged a sCMOS camera (pco.edge, PCO AG, Kelheim, Germany) mounted on a Axio Observer D1 (Jena, Germany) microscope. The pore dynamics were analyzed using an automated approach as in (81), which considers the slow regime of pore closure(41, 58).

### Cell handling and liposome incubation

Human embryonic kidney (HEK) cells (American-type culture collection, no. CRL-1573, lot no. 63 966 486) were used for the cell experiments as in our recent study(62). Dulbecco’s minimal essential medium (DMEM, Giboc), fetal bovine serum (Gibco), phosphate buffered saline (PBS, Gibco), trypsin (0.05%, Gibco), trypan blue (0.4%, Sigma-Aldrich), a μ-slide 8 well chambers (Ibidi), 35 mm petri dishes (Grenier Bio-One), and a Neubauer hemacytometer (Brand). HEK cells were cultured in DMEM medium supplemented with 10% serum (complete medium) at 37 °C under 5% CO_2_ in a humidified atmosphere (All results are from cell cultures that tested negative for mycoplasma). Cells were seeded into μ-slide 8 well dishes one day before the experiments. For confocal imaging, the cells were moved to the microscope stage at 37 °C and the cells were washed once with a sucrose 300 mM solution (isotonic to culture media) to remove the serum. In the control permeability experiment (no liposomes), the culture medium contained 10 μM SRB and 0.1 mg/ml Dextran 10 kDa. For incubation with liposomes, a liposome solution (500 μM, lipid concentration) in sucrose 300 mM, prewarmed at 37 °C, was added to the cells. Cells were images either after 30 minutes incubation or imaged for 30 minutes followed liposome addition.

### Microscopy imaging

Confocal fluorescence microscopy was performed on a Zeiss LSM 710 scanning confocal microscope. Images were acquired through a water immersion C-Apochromat 40X/1.20 W Korr M27 objective. The spatial and temporal resolutions used were adjusted according to the samples; in general imaging size of 212.55 μm x 212.55 μm (1024 × 1024 pixels) was used in a frame scanning mode in a singular direction, with a pixel size of 0.21 mm with 4 line averaging and a bit depth of 8. The green dyes Bodipy C_16_, Dextran-FITC 3 kDa and Dextran 10 kDa were excited using a 488 nm argon laser, and emission was detected between 495 and 555 nm. The orange dyes DPPE-Rh and SRB were excited with a 543 nm HeNe laser line and emission was detected in the range between 555 and 600 nm. The far-red dye Atto-647N was excited with a 633 nm HeNe laser line and its emission was detected in the range between 640 and 800 nm. To minimize crosstalk between channels, the images were scanned in the sequential mode.

Fluorescence lifetime imaging microscopy (FLIM) experiments were carried out as in(89). In short, images were acquired on a Microtime200 microscope (PicoQuant) built on an inverted microscope (Olympus IX73) equipped with time correlated single photon counting (TCSPC). The samples were imaged through a 100× (1.4 NA) oil immersion objective (UPLSAPO, Olympus). Bodipy C_16_ was excited using a 481 nm laser and its emission was collected using a 525/50 nm band pass filter. The images were acquired using the SymPhoTime 64 software and all samples were excited with a pulsed 20 MHz repetition rate. The samples were imaged with a 128 × 128 pixels, 1 ms dwell-time and ≈300 μm/pixels with typical acquisition times of ≈27 s. For analysis, the signal on the GUV membrane at the equator was manually selected for analysis and all pixels binned for fitting. Bodipy C_16_ fluorescence decays were fitted using an n-exponential tail fit with a single- (in the absence of an acceptor) or bi-exponential decay model (in the presence of the DPPE-Rh FRET acceptor):

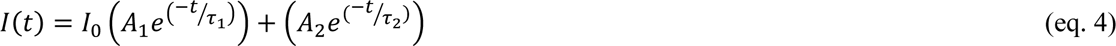

*I*(*t*) is the intensity at time *t* and *I*_0_ is the intensity at *t* = 0. *A*_1_ and *A*_2_ are pre-exponentials factors associated with lifetime components *τ*_1_ and *τ*_2_, respectively. The amplitude-weighted mean fluorescence lifetime in the presence of the FRET acceptor (*τ*) could be calculated as

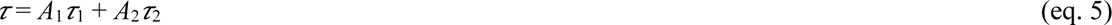

The degree-of-filling method measures the equilibration of a fluorescent solute between the vesicle interior and the external solution(51), and it was used to assess GUV permeability upon fusion with the LUVs. The GUVs were incubated with a given LUV concentration for 10 minutes in the presence of a specific fluorescent leakage reporter, after which they were diluted ∼ 3x (the lower LUV concentration) in a solution containing the identical concentration of the reporter. The degree of filling was calculated as

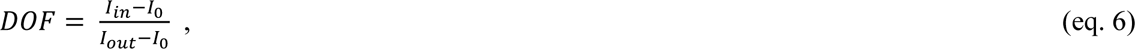

where I_in_ and I_out_ are the fluorescence intensity measured inside and outside the GUV and I_0_ is the background fluorescence. Values range from 0 (non-permeable) to 1 (permeable). To account for background noise, we define 0.1 as a permeability threshold(49, 51).

The FRET efficiency (*E*_*FRET*_) was calculated by measuring the fluorescence lifetime of the FRET donor (Bodipy C_16_) on individual GUVs (illustrated in Figure S1A) as E_FRET_ = 1-τ_LUV_/τ_control_(90). Here, τ_LUV_ and τ_control_ represent the control fluorescence lifetime in the presence and absence of LUVs, respectively. We produced a standard calibration curve by measuring *E*_*FRET*_ on several GUVs for increasing molar fraction of the FRET acceptor (DPPE-Rh) while keeping the donor (Bodipy C_16_) at constant molar fraction (0.5 mol%) (Figure S1), which allows us to obtain the ratio acceptor/donor from E_FRET_ values(19). In order to obtain this analytically, we fitted E_FRET_ as function of acceptor/donor using a Hill equation:

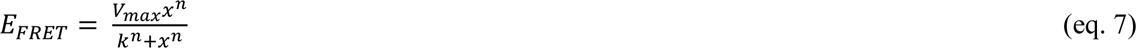

where *x* represents the acceptor/donor ratio, and *V*_*max*_, *n* and *k* are fitting parameters without a defined physical meaning for our purpose. Inversion of eq. 2 then gives us an analytical expression to correlate

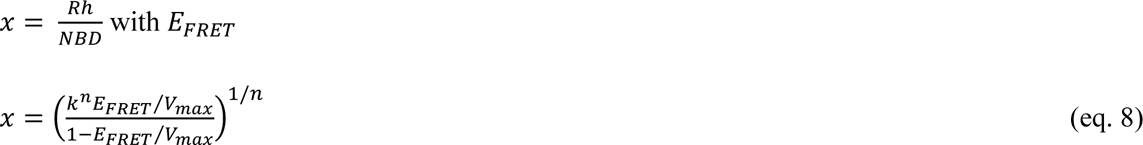

The parameters used to fit the calibration curve (Figure S1) were *V*_*max*_ = 0.9, *n* = 2.2 and *k* = 5.4. Unlike intensity-based measurements, where the calibration will change depending on experimental conditions (i.e. microscope settings, such as laser intensity, spectral range), the FLIM-FRET calibration curve provides absolute values at a given fraction of donor, and hence the obtained fitting parameters will not depend on experimental factors, hence being more robust.

From the acceptor/donor ratio measured in the LUV-GUV fusion experiments, and Bodipy C_16_ molar fraction 0.5 mol%, we can obtain the molar fraction of the acceptor *X*_*Rh*_ transferred to the GUV via fusion (considering that the composition of the LUVs is DOPE:DOTAP:DPPE-Rh 10:10:1), i.e. *X*_*fuso*_= 21 *X*_*Rh*_. We emphasize that the analysis does not account for the dilution of the FRET donor probe molecules in the GUV membrane due to fusion, yielding somewhat overestimated FRET values. Next, we calculate the number of LUVs (*n*_*LUV*_) fused with the GUV from the ratio between the area increment brought by the LUVs. Because *X*_*fuso*_is the fraction of lipids coming from the LUVs, then 1 − *X*_*fuso*_corresponds to the fraction of original lipids in the GUV, and *X*_*fuso*_⁄(1 − *X*_*fuso*_) is the relative increment in lipid number in the GUV upon fusion of fusogenic LUVs. Assuming that the area increment of the GUVs is proportional to the number of lipids gained via fusion - lipids have roughly the same area per molecule

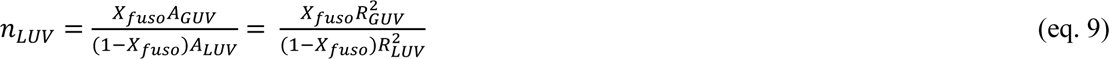

where *R*_*GUV*_ is the initial radius of the particular GUV and *R*_*LUV*_ the mean radius of LUVs, measured with DLS (*R*_*LUV*_ = 60 nm)(91).

The lipid number asymmetry N_out_/N_in_ can be calculated as the ratio between the external and internal area, i.e. (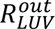/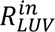)^2^. Assuming that the size of the LUV is given by its external radius, 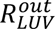 ≡ *R*_*LUV*_= 60 nm, then the internal radius will be this value minus the bilayer thickness (∼5 nm), 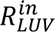 ≅ 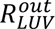 − 5 = 55 nm. In other words, each LUV has a lipid asymmetry described by the lipid ratio (*N_out_*/*N_in_*)*_LUV_* = (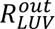/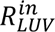)^2^ ≅ 1.2 (i.e. 55% of the lipids reside in the outer and 45% in the inner leaflet), and hence asymmetry is given by

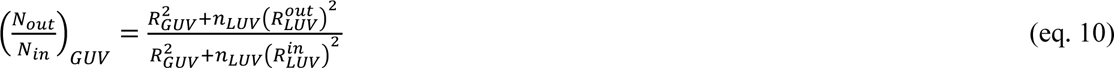

Edge tension experiments were performed as in(81). The GUVs were diluted ∼10× in isotonic glucose and placed in an electrofusion chamber containing two parallel cylinder electrodes (92 μm radius spaced by 500 μm (43)). The chamber was connected to an Eppendorf multiporator (Eppendorf), wherein pulse strength and duration can be controlled from 50 to 300 V and 50 to 300 μs, respectively. Imaging was performed on a Zeiss Axiovert 200 (Jena) phase contrast microscope equipped with an sCMOS camera (pco.edge 4.2, PCO AG) for fast recordings (300 frames per second, fps used here), through a 63x magnification air objective (NA 0.75). Field strength of 150 V (3 kV/cm−1) and 150 μs duration pulses were applied, and pore closure dynamics were analysed by tracking pore sizes using a freely available Python-based software (81). For a circular pore of r radius,

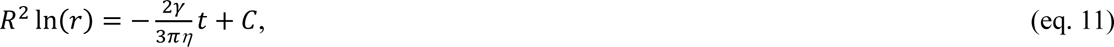

where η = 1.33 cP is the viscosity of the aqueous medium) and C is a time-independent constant that varies per experiment(41, 92). By plotting the pore closure *R*^2^ln (*r*/*l*) versus time, where R is the GUV radius and the reference length *l*=1 makes the logarithm dimensionless, γ can be directly calculated from the slope *a* of the linear regime of slow pore closure given by

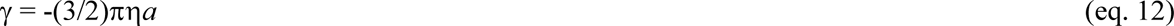

### Coarse grained molecular dynamics simulations

All simulations were executed with GROMACS version 2022.1(93)using the Martini 3 CG forcefield(94, 95) All CG membrane systems used in this study were constructed using TS2CG(96), an MD tool designed to generate initial configurations for lipid membrane structures with user-defined compositions. For all simulations in this study an initial system size of 40×40×20 nm was used. The details and membrane composition of the performed experiments can be found in Table S1. After generating the initial configuration an initial energy minimization was performed using a steepest descent algorithm. This was followed by three equilibration runs with integration steps of 0.02, 0.2 and 2 fs, totalling 10 ns. To simulate pore nucleation, a pore was artificially created using a cylindrical restraint that gradually increased in diameter over the course of 20 ns. Simulation parameters were based on standard values used in the Martini benchmarking paper(97). The fixed area simulations were performed in the NPT ensemble at 310 K. A pressure of 1 bar was applied in the z-direction, while in the lateral directions the system was not pressure coupled; instead, the compressibility modulus was set to zero. Both pressure and the temperature were maintained using an anisotropic Berendsen coupling(98) and velocity rescaling thermostat(99), respectively. Furthermore, an integration step of 20 fs was chosen for the production runs using a leapfrog algorithm for integrating the equations of motion. VMD (100) was used for rendering the snapshots of the simulations. For calculating the membrane properties during the simulation, in-house Python scripts were used.

## Supporting information

Supplementary results

## Author Contributions

RBL designed the study, performed most experiments, analysed the data and wrote the first draft of the manuscript. MvT performed the simulations. RRMC performed the edge tension experiments. MK aided in the design of the simulations. CJR performed part of the cell experiments. KAR supervised the edge tension experiments. SJM and WHR supervised the study. All authors read and edited the manuscript and approved the final version.

## Acknowledgements

The authors thank Melanie König for useful discussions in the early stages of the project. SJM acknowledges funding from the ERC with the Advanced grant “COMP-O-CELL”.

## REFERENCES

1. R. Whittam & K. P. Wheeler, TRANSPORT ACROSS CELL MEMBRANES. Annu. Rev. 32, 21–60 (2026).

2. D. Drew, O. Boudker, Ion and lipid orchestration of secondary active transport. Nature 626, 963–974 (2024).

3. T. Harayama, H. Riezman, Understanding the diversity of membrane lipid composition. Nat. Rev. Mol. Cell Biol., 281–296 (2018).

4. K. Simons, M. J. Gerl, Revitalizing membrane rafts: New tools and insights. Nat. Rev. Mol. Cell Biol. 11, 688–699 (2010).

5. E. Sezgin, I. Levental, S. Mayor, C. Eggeling, The mystery of membrane organization: Composition, regulation and roles of lipid rafts. Nat. Rev. Mol. Cell Biol. 18, 361–374, (2017).

6. J. E. Rothman, J. Lenard, Membrane Asymmetry The nature of membrane asymmetry provides clues to the puzzle of how membranes are assembled. Science 195(4280), 743–753 (1977).

7. M. Doktorova, I. Levental, F. A. Heberle, Seeing the Membrane from Both Sides Now: Lipid Asymmetry and Its Strange Consequences. Cold Spring Harb. Perspect. Biol. 15 (2023).

8. M. Doktorova, J. L. Symons, I. Levental, Structural and functional consequences of reversible lipid asymmetry in living membranes. Nat. Chem. Biol. 16, 1321–1330 (2020).

9. J. H. Lorent, et al. Plasma membranes are asymmetric in lipid unsaturation, packing and protein shape. Nat. Chem. Biol. 16, 644–652 (2020).

10. S. L. Liu, et al. Orthogonal lipid sensors identify transbilayer asymmetry of plasma membrane cholesterol. Nat. Chem. Biol. 13, 268–274 (2017).

11. M. Aghaaminiha, A. M. Farnoud, S. Sharma, Quantitative relationship between cholesterol distribution and ordering of lipids in asymmetric lipid bilayers. Soft Matter 17, 2742–2752 (2021).

12. M. Varma, M. Deserno, Distribution of cholesterol in asymmetric membranes driven by composition and differential stress. Biophys. J. 121, 4001–4018 (2022).

13. P. F. Devaux. Static and Dynamic Lipid Asymmetry in Cell Membranes. Biochemistry 30(5), 1163–1173 (1991).

14. M. Deserno, Biomembranes balance many types of leaflet asymmetries. Curr. Opin. Struct. Biol. 87:102832 (2024).

15. G. J. Schütz, G. Pabst, The asymmetric plasma membrane—A composite material combining different functionalities?: Balancing Barrier Function and Fluidity for Effective Signaling. BioEssays 45(12), 2300116 (2023).

16. M. Doktorova, et al. Cell membranes sustain phospholipid imbalance via cholesterol asymmetry. Cell 188, 2586–2602.e24 (2025).

17. S. Sanyal, A. K. Menon, Flipping lipids: Why an’ what’s the reason for? ACS Chem. Biol. 4(11), 895–909 (2009).

18. T. H. Ogunmowo, et al. Membrane compression by synaptic vesicle exocytosis triggers ultrafast endocytosis. Nat. Commun. 14, 2888 (2023).

19. R. B. Lira, R. M. Cavalcanti, K. A. Riske, R. Dimova, The Hidden Cost of Fusion: Intrinsic Asymmetry in Vesicle Fusion Restricts Synthetic Cell Growth. (2025). doi: 10.1101/2025.09.22.677743.

20. M. S. Miettinen, R. Lipowsky, Bilayer Membranes with Frequent Flip-Flops Have Tensionless Leaflets. Nano Lett. 19, 5011–5016 (2019).

21. A. Sreekumari, R. Lipowsky, Large stress asymmetries of lipid bilayers and nanovesicles generate lipid flip-flops and bilayer instabilities. Soft Matter 18, 6066–6078 (2022).

22. M. P. Sheetz, S. J. Singert, “Biological Membranes as Bilayer Couples. A Molecular Mechanism of Drug-Erythrocyte Interactions. PNAS 71, (11) 4457–4461 (1974).

23. E. Farge, P. F. Devaux, Shape changes of giant liposomes induced by an asymmetric transmembrane distribution of phospholipids. Biophys. J. 61, 347–357 (1992).

24. D. Blanken, D. Foschepoth, A. C. Serrão, C. Danelon, Genetically controlled membrane synthesis in liposomes. Nat. Commun. 11, 4317 (2020).

25. T. F. Zhu, J. W. Szostak, Coupled growth and division of model protocell membranes. J. Am. Chem. Soc. 131, 5705–5713 (2009).

26. I. A. Chen, J. W. Szostak, A kinetic study of the growth of fatty acid vesicles. Biophys. J. 87, 988–998 (2004).

27. R. B. Lira, T. Robinson, R. Dimova, K. A. Riske, Highly Efficient Protein-free Membrane Fusion: A Giant Vesicle Study. Biophys. J. 116, 79–91 (2019).

28. R. B. Lira, R. Dimova, Fusion assays for model membranes: a critical review. Advances in Biomembranes and Lipid Self-Assembly 30, 229–270 (2019).

29. M. P. K. Frewein, et al. Distributing aminophospholipids asymmetrically across leaflets causes anomalous membrane stiffening. Biophys. J. 122, 2445–2455 (2023).

30. Y. Elani, et al. Measurements of the effect of membrane asymmetry on the mechanical properties of lipid bilayers. Chemical Communications 51, 6976–6979 (2015).

31. F. S. C. Leomil, M. Stephan, S. Pramanik, K. A. Riske, R. Dimova, Bilayer Charge Asymmetry and Oil Residues Destabilize Membranes upon Poration. Langmuir 40, 4719–4731 (2024).

32. S. Esteban-Martín, H. Jelger Risselada, J. Salgado, S. J. Marrink, Stability of asymmetric lipid bilayers assessed by molecular dynamics simulations. J. Am. Chem. Soc. 131, 15194–15202 (2009).

33. M. Aleksanyan, R. B. Lira, J. Steinkühler, R. Dimova, GM1 asymmetry in the membrane stabilizes pores. Biophys. J. 121, 3295–3302 (2022).

34. A. A. Gurtovenko, I. Vattulainen, Molecular mechanism for lipid flip-flops. Journal of Physical Chemistry B 111, 13554–13559 (2007).

35. J. S. Allhusen, J. C. Conboy, The ins and outs of lipid flip-flop. Acc. Chem. Res. 50, 58–65 (2017).

36. J. D. Litster, Stability of lipid bilayers and red blood cell membranes. Physics Letters A, 53(3), 193–194 (1975).

37. E. Evans, D. Needham, “Physical Properties of Surfactant Bilayer Membranes: Thermal Transitions, Elasticity, Rigidity, Cohesion, and Colloidal Interactions” J. Phys. Chem. 91, 4219–4228 (1987).

38. S. A. Akimov, et al. Pore formation in lipid membrane I: Continuous reversible trajectory from intact bilayer through hydrophobic defect to transversal pore. Sci. Rep. 7 (2017).

39. R. Lipowsky, The many faces of membrane tension for biomembranes and vesicles. Faraday Discuss. 259, 234–263 (2025).

40. W. Rawicz, B. A. Smith, T. J. McIntosh, S. A. Simon, E. Evans, Elasticity, strength, and water permeability of bilayers that contain raft microdomain-forming lipids. Biophys. J. 94, 4725–4736 (2008).

41. T. Portet, R. Dimova, A new method for measuring edge tensions and stability of lipid bilayers: Effect of membrane composition. Biophys. J. 99, 3264–3273 (2010).

42. B. Mattei, R. B. Lira, K. R. Perez, K. A. Riske, Membrane permeabilization induced by Triton X-100: The role of membrane phase state and edge tension. Chem. Phys. Lipids 202, 28–37 (2017).

43. K. A. Riske, R. Dimova, Electro-deformation and poration of giant vesicles viewed with high temporal resolution. Biophys. J. 88, 1143–1155 (2005).

44. R. B. Lira, R. Dimova, K. A. Riske, Giant unilamellar vesicles formed by hybrid films of agarose and lipids display altered mechanical properties. Biophys. J. 107, 1609–1619 (2014).

45. T. Sakuragi, S. Nagata, Regulation of phospholipid distribution in the lipid bilayer by flippases and scramblases. Nat. Rev. Mol. Cell Biol. 24, 576–596 (2023).

46. S. Sanyal, A. K. Menon, Flipping lipids: Why an’ what’s the reason for? ACS Chem. Biol. (2009).

47. R. Watanabe, T. Sakuragi, H. Noji, S. Nagata, Single-molecule analysis of phospholipid scrambling by TMEM16F. Proc. Natl. Acad. Sci. U. S. A. 115, 3066–3071 (2018).

48. R. Lipowsky, Spontaneous tubulation of membranes and vesicles reveals membrane tension generated by spontaneous curvature. Faraday Discuss. 161, 305–331 (2012).

49. R. B. Lira, et al. The underlying mechanical properties of membranes tune their ability to fuse. Journal of Biological Chemistry 299(12), 105430 (2023).

50. R. R. M. Cavalcanti, R. B. Lira, E. J. Ewins, R. Dimova, K. A. Riske, Efficient liposome fusion to phase-separated giant vesicles. Biophys. J. 122, 2099–2111 (2023).

51. S. Bleicken, O. Landeta, A. Landajuela, G. Basañez, A. J. García-Sáez, Proapoptotic Bax and Bak proteins form stable protein-permeable pores of tunable size. Journal of Biological Chemistry 288, 33241–33252 (2013).

52. U. Ros, L. Pedrera, A. J. Garcia-Saez, Techniques for studying membrane pores. Curr. Opin. Struct. Biol. 69, 108–116 (2021).

53. J. K. Armstrong, R. B. Wenby, H. J. Meiselman, T. C. Fisher, The hydrodynamic radii of macromolecules and their effect on red blood cell aggregation. Biophys. J. 87, 4259–4270 (2004).

54. J. Hammond, et al. Membrane Fusion-Based Drug Delivery Liposomes Transiently Modify the Material Properties of Synthetic and Biological Membranes. Small 21(12), 2408039 (2025).

55. H. A. Faizi, S. L. Frey, J. Steinkühler, R. Dimova, P. M. Vlahovska, Bending rigidity of charged lipid bilayer membranes. Soft Matter 15, 6006–6013 (2019).

56. K. Olbrich, W. Rawicz, D. Needham, E. Evans, Water permeability and mechanical strength of polyunsaturated lipid bilayers. Biophys. J. 79, 321–327 (2000).

57. P. H. Puech, N. Borghi, E. Karatekin, F. Brochard-Wyart, Line Thermodynamics: Adsorption at a Membrane Edge. Phys. Rev. Lett. 90, 4 (2003).

58. E. Karatekin, et al. Cascades of transient pores in giant vesicles: Line tension and transport. Biophys. J. 84, 1734–1749 (2003).

59. R. B. Lira, F. S. C. Leomil, R. J. Melo, K. A. Riske, R. Dimova, To Close or to Collapse: The Role of Charges on Membrane Stability upon Pore Formation. Advanced Science 8 (2021).

60. V. K. Malik, O. S. Pak, J. Feng, Curvature-Assisted Vesicle Explosion Under Light-Induced Asymmetric Oxidation. Advanced Science 11(9), 2004068 (2024).

61. H. Pera, J. M. Kleijn, F. A. M. Leermakers, On the edge energy of lipid membranes and the thermodynamic stability of pores. Journal of Chemical Physics 142, 034101 (2015).

62. J. Hammond, et al. Membrane Fusion-Based Drug Delivery Liposomes Transiently Modify the Material Properties of Synthetic and Biological Membranes. Small (2025). 10.1002/smll.202408039.

63. R. B. Lira, J. Willersinn, B. V. K. J. Schmidt, R. Dimova, Selective partitioning of (Biomacro)molecules in the crowded environment of double-hydrophilic block copolymers. Macromolecules 53, 10179–10188 (2020).

64. D. P. Tieleman, S. J. Marrink, Lipids out of equilibrium: Energetics of desorption and pore mediated flip-flop. J. Am. Chem. Soc. 128, 12462–12467 (2006).

65. J. Solon, et al. Negative tension induced by lipid uptake. Phys. Rev. Lett. 97, 098103 (2006).

66. M. Yu, R. B. Lira, K. A. Riske, R. Dimova, H. Lin, Ellipsoidal Relaxation of Deformed Vesicles. Phys. Rev. Lett. 115, 128303 (2015).

67. N. C. Gauthier, T. A. Masters, M. P. Sheetz, Mechanical feedback between membrane tension and dynamics. Trends Cell Biol. 22(10), 527–535 (2012).

68. G. Saulis, M. Venslauskas, J. Naktinis. Kinetics of pore resealing in cell membranes after electroporation. Journal of Electroanalytical Chemistry and Interfacial Electrochemistry 321(1), 1–13 (1991).

69. A. G. Pakhomov, et al. Multiple nanosecond electric pulses increase the number but not the size of long-lived nanopores in the cell membrane. Biochim. Biophys. Acta Biomembr. 1848, 958–966 (2015).

70. 42. R. B. Lira, and W. H. Roos. 2026. Influence of substrate on supported lipid bilayers: Membrane adhesion, stretching, pores, and remodeling. Langmuir 42:5985–5999.

71. T. Pomorski, A. K. Menon, Lipid flippases and their biological functions. Cellular and Molecular Life Sciences 63, 2908–2921 (2006).

72. R. B. Lira, C. Dekker, Lipid flip flop regulates the shape of growing and dividing synthetic cells. (2025). http://biorxiv.org/lookup/doi/10.1101/2025.04.23.650179.

73. M. A. Goren, et al. Constitutive phospholipid scramblase activity of a G protein-coupled receptor. Nat. Commun. 5, 5115 (2014).

74. H. Jahn, et al. Phospholipids are imported into mitochondria by VDAC, a dimeric beta barrel scramblase. Nat. Commun. 14, 8115 (2023).

75. J. A. Coleman, A. L. Vestergaard, R. S. Molday, B. Vilsen, J. P. Andersen, Critical role of a transmembrane lysine in aminophospholipid transport by mammalian photoreceptor P 4-ATPase ATP8A2. PNAS 109, 1449–1454 (2012).

76. I. Menon, et al. Opsin is a phospholipid flippase. Current Biology 21, 149–153 (2011).

77. A. Papadopulos, et al. Flippase activity detected with unlabeled lipids by shape changes of giant unilamellar vesicles. Journal of Biological Chemistry 282, 15559–15568 (2007).

78. L. J. Starke, C. Allolio, J. S. Hub, How pore formation in complex biological membranes is governed by lipid composition, mechanics, and lateral sorting. PNAS Nexus 4(3), (2025).

79. E. A. Cino, D. P. Tieleman, Curvature-based sorting of eight lipid types in asymmetric buckled plasma membrane models. Biophys. J. 121, 2060–2068 (2022).

80. J. Pan, et al. Molecular structures of fluid phase phosphatidylglycerol bilayers as determined by small angle neutron and X-ray scattering. Biochim. Biophys. Acta Biomembr. 1818, 2135–2148 (2012).

81. F. S. C. Leomil, M. Zoccoler, R. Dimova, K. A. Riske, A. Bateman, PoET: automated approach for measuring pore edge tension in giant unilamellar vesicles. Bioinformatics Advances, 1(1), vbab037 (2021).

82. S. Chakraborty, et al. How cholesterol stiffens unsaturated lipid membranes. PNAS 117 (36) 21896–21905 (2020).

83. Gerrit van Meer, Dennis R. Voelker & Gerald W. Feigenson. Membrane lipids: where they are and how they behave. Nature Reviews Molecular Cell Biology 9, 112–124 (2008).

84. Miao, L.; Seifert, U.; Wortis, M.; and Döbereiner, H.-G. Budding Transitions of Fluid-Bilayer Vesicles: The Effect of Area-Difference Elasticity. Phys. Rev. E, 49, 5389–5407 (1994).

85. Jarić, M.; Seifert, U.; Wintz, W.; and Wortis, M. Vesicular Instabilities: The Prolate-to-Oblate Transition and Other Shape Instabilities of Fluid Bilayer Membranes. Phys. Rev. E 52, 6623–6634 (1995).

86. A. Weinberger, et al. Gel-assisted formation of giant unilamellar vesicles. Biophys. J. 105, 154–164 (2013).

87. R. B. Lira, et al. Studies on intracellular delivery of carboxyl-coated CdTe quantum dots mediated by fusogenic liposomes. J. Mater. Chem. B 1, 4297–4305 (2013).

88. S. Aranda, K. A. Riske, R. Lipowsky, R. Dimova, Morphological transitions of vesicles induced by alternating electric fields. Biophys. J. 95 (2008).

89. R. B. Lira, et al. Fluorescence lifetime imaging microscopy of flexible and rigid dyes probes the biophysical properties of synthetic and biological membranes. Biophys. J. 123, 1592–1609 (2024).

90. J. R. Lakowicz, “Principles of Fluorescence Spectroscopy Third Edition.”

91. R. R. M. Cavalcanti, R. B. Lira, K. A. Riske, Membrane Fusion Biophysical Analysis of Fusogenic Liposomes. Langmuir 38, 10430–10441 (2022).

92. F. Brochard-Wyart, P. G. De Gennes, O. Sandre, Transient pores in stretched vesicles: role of leak-out. Physica A: Statistical Mechanics and its Applications 278, 32–51 (2000).

93. M. J. Abraham, et al. Gromacs: High performance molecular simulations through multi-level parallelism from laptops to supercomputers. SoftwareX 1–2, 19–25 (2015).

94. K. B. Pedersen, et al. The Martini 3 Lipidome: Expanded and Refined Parameters Improve Lipid Phase Behavior. ACS Cent. Sci. 11, 1598–1610 (2025).

95. L. Borges-Araújo, et al. Martini 3 Coarse-Grained Force Field for Cholesterol. J. Chem. Theory Comput. 19, 7387–7404 (2023).

96. W. Pezeshkian, M. König, T. A. Wassenaar, S. J. Marrink, Backmapping triangulated surfaces to coarse-grained membrane models. Nat. Commun. 11, 2296 (2020).

97. D. H. De Jong, S. Baoukina, H. I. Ingólfsson, S. J. Marrink, Martini straight: Boosting performance using a shorter cutoff and GPUs. Comput. Phys. Commun. 199, 1–7 (2016).

98. H. J. C. Berendsen, J. P. M. Postma, W. F. Van Gunsteren, A. Dinola, J. R. Haak, Molecular dynamics with coupling to an external bath. J. Chem. Phys. 81, 3684–3690 (1984).

99. G. Bussi, D. Donadio, M. Parrinello, Canonical sampling through velocity rescaling. Journal of Chemical Physics 126, 014101(2007).

100. W. Humphrey, A. Dalke, K. Schulten, VMD: Visual Molecular Dynamics. Journal of Molecular Graphics 14(1), 33–38 (1996).

