## Supplementary results for "MEMBRANE PORES ACT AS SELF-RESEALING LIPID SCRAMBLASES"

to the paper


### SI Figures

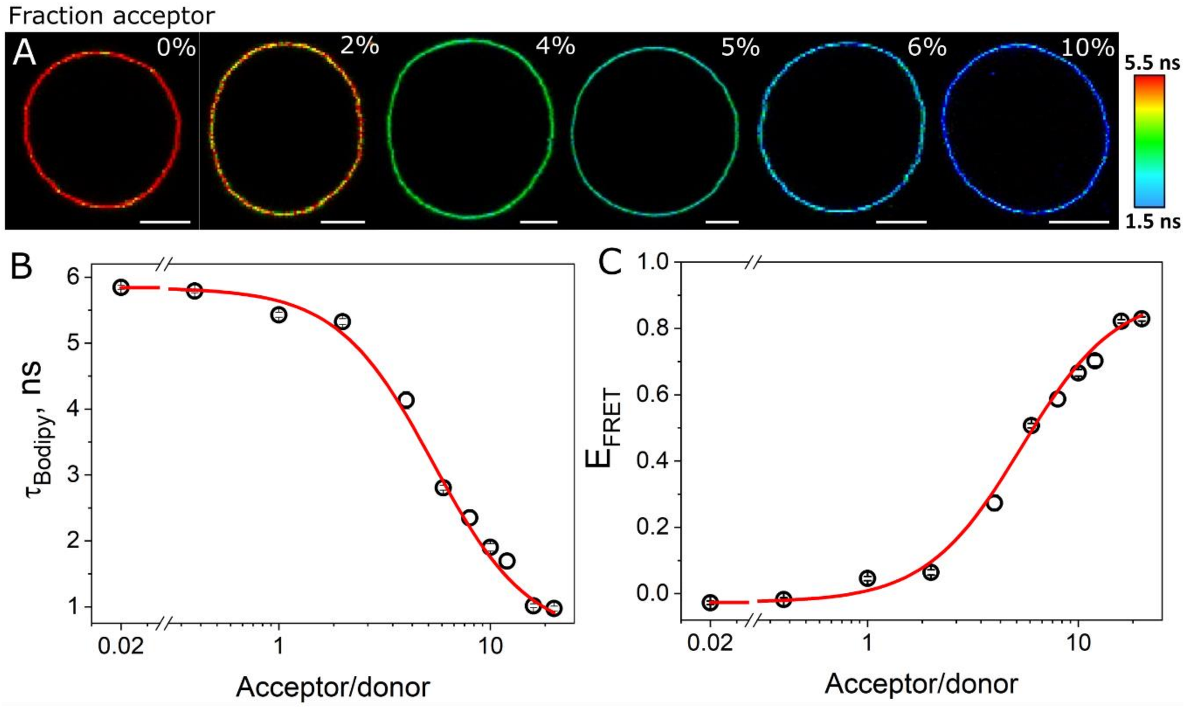


**Figure S1**. FLIM-FRET standard calibration curve. A, representative DOPC GUVs containing 0.5 mol% Bodipy $C_{16}$ as a FRET donor, and increasing fractions of DPPE-Rh (indicated in the respective snapshots) as a FRET acceptor. Bodipy $C_{16}$ lifetime map is shown on the right. Scale bars: 10 µm. The FRET donor Bodipy $C_{16}$ was chosen for its insensitivity to environmental changes[1] (i.e. lipid composition), thereby its lifetime changes report exclusively FRET changes. B and C, measured Bodipy $C_{16}$ lifetime, and calculated FRET efficiency ($E_{FRET}$) as a function of acceptor/donor ratio. Data points represent mean and s.d from > 15 individual GUVs. The lines are fits to equations 7 in the main text.

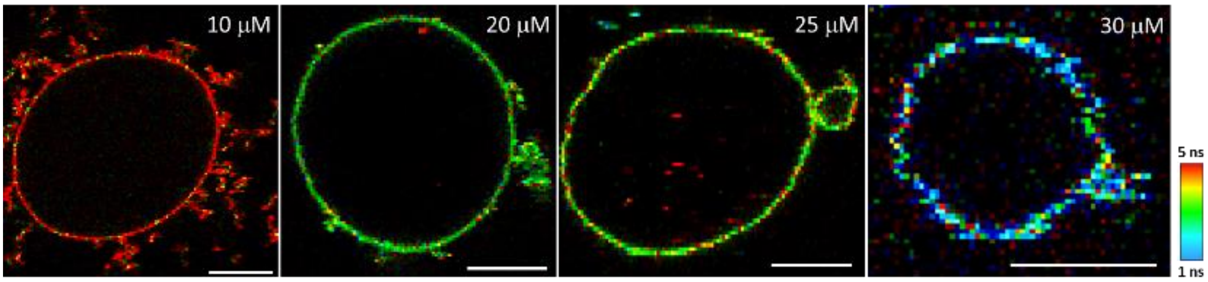


**Figure S2.** GUV budding upon fusion with LUVs. Representative GUVs made of equimolar fractions of DOPC:DOPG:DOPE:chol and labelled with 0.5 mol% Bodipy $C_{16}$ FRET donor were incubated with increasing concentrations (indicated in the snapshots) of LUVs labelled with 2 mol% DPPE-Rh FRET acceptor. Changes in colour represent reduction in donor lifetime. Note outward budding whose lifetime is identical to that in the flat membrane regions. Scale bars: 7 µm.

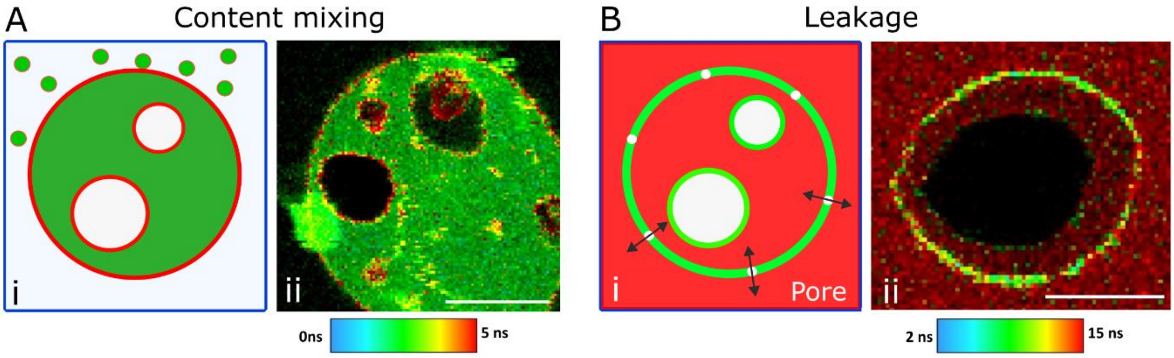


**Figure S3.** GUV Content mixing and leakage upon fusion with LUVs. A, sketch of LUV fusion that results in mixing of the vesicle contents (green) upon full-fusion with a GUV and that contains smaller vesicles in its interior (i). On the right (ii), representative content mixing experiment of non-labelled LUVs made of entrapping 0.1 mg/ml Dextran 10 kDa. Note that content transfer is limited to the fusing GUV, whereas internal vesicles remain devoid of the probe. B, sketch of fusion-mediated leakage of a GUV containing smaller vesicles in its interior in the presence of a leakage marker (red) (i). On the right (ii), representative leakage of KU530 in the GUV in direct contact with the LUVs. Note that the internal vesicle remains intact. LUV composition: DOTAP:DOPE (5:5 molar ratio). GUV composition: DOPC:$_B$PS (50:50, molar ratio) GUV, labelled with 0.5 mol% Bodipy $C_{16}$. Scale bars: 10 µm.

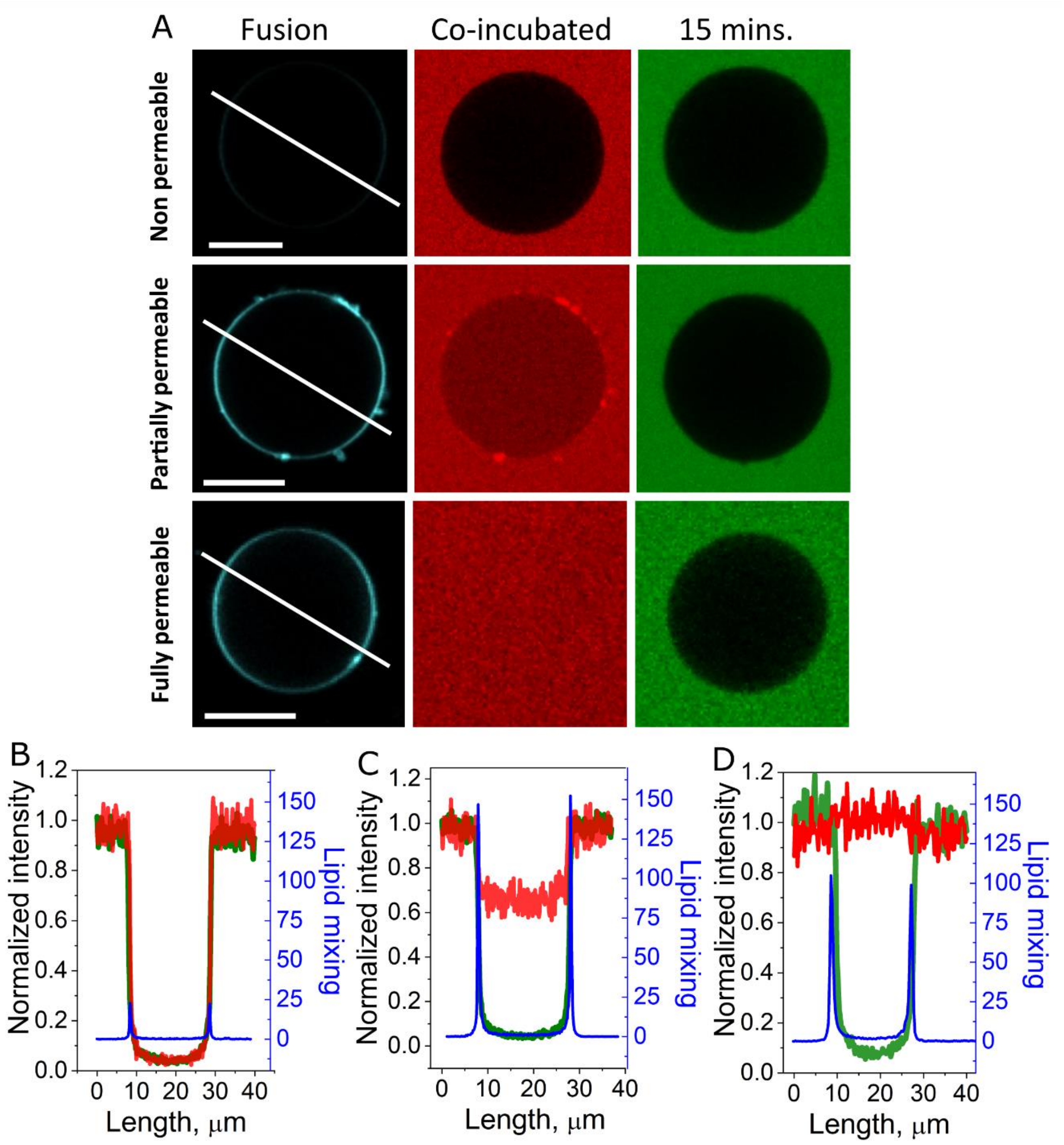


**Figure S4.** Leakage pores reseal. A, representative equimolar DOPC:DOPG:DOPE:Chol GUVs incubated with 15 μM (lipid concentration) DOTAP:DOPE LUVs labelled with DOPE-Atto647N (cyan) and co-incubated with 20 μM SRB (red) used as a leakage marker. After ~ 25 minutes, 10 μM calcein (green) is added to the samples to assess for open/closed GUV pores. Note the different levels of SRB permeation. Importantly, at later points, the GUVs are not permeable to calcein, indicating resealing of leakage pores. Scale bars: 10 μm. B-D, intensity traces along the white line shown in A for the three fluorescence channels for the non permeable, partially permeable and fully permeable GUVs, respectively.

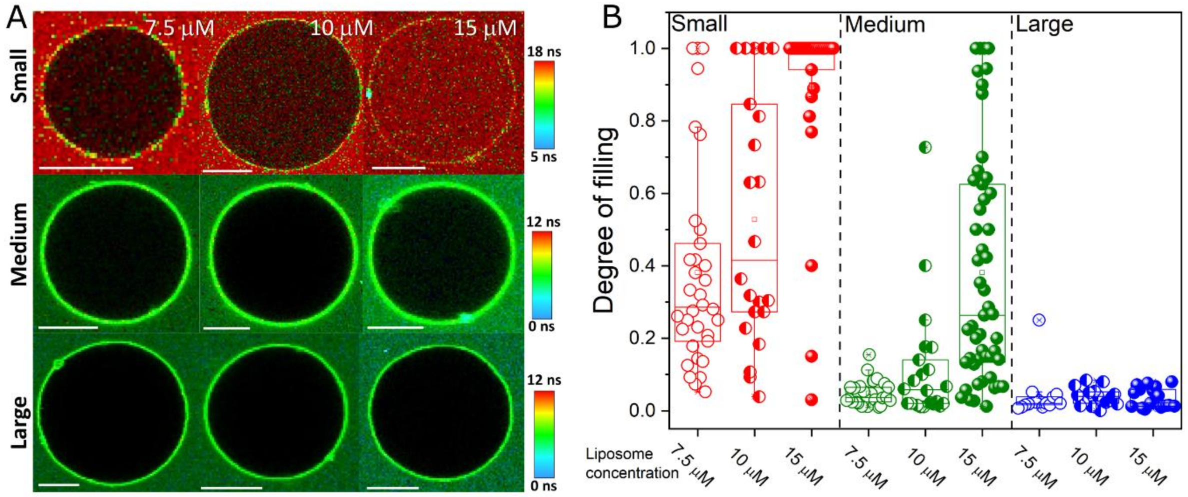


**Figure S5.** Increasing LUV fusion results in larger leakage pores. A, representative FLIM snaphots of equimolar DOPC:DOPG:Chol:DOPE GUVs labelled with 0.5 mol% Bodipy $C_{12}$ incubated with increasing concentrations (indicated in the images) of LUVs (unlabelled DOTAP:DOPE) in the presence of the small probe KU530, the medium probe Dextran 3 kDa, and the large probe Dextran 10 kDa. Scale bars: 10 µm. B, degree of filling measurements for many GUVs for increasing LUV concentrations and co-incubated with the probes of increasing sizes.

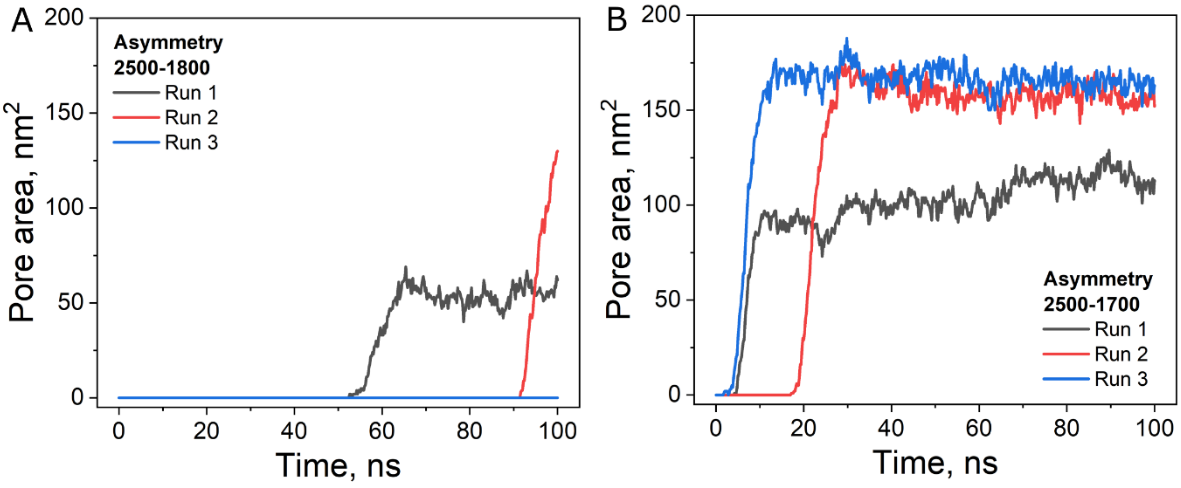


**Figure S6**. Stochastic pore opening. Two representative asymmetries. Note the differences in pore area and onset of pore formation for different runs at these two intermediate asymmetries. While for lower asymmetries, pores do not spontaneously open, for higher asymmetries, different runs give more similar values for area and onset of pore formation.

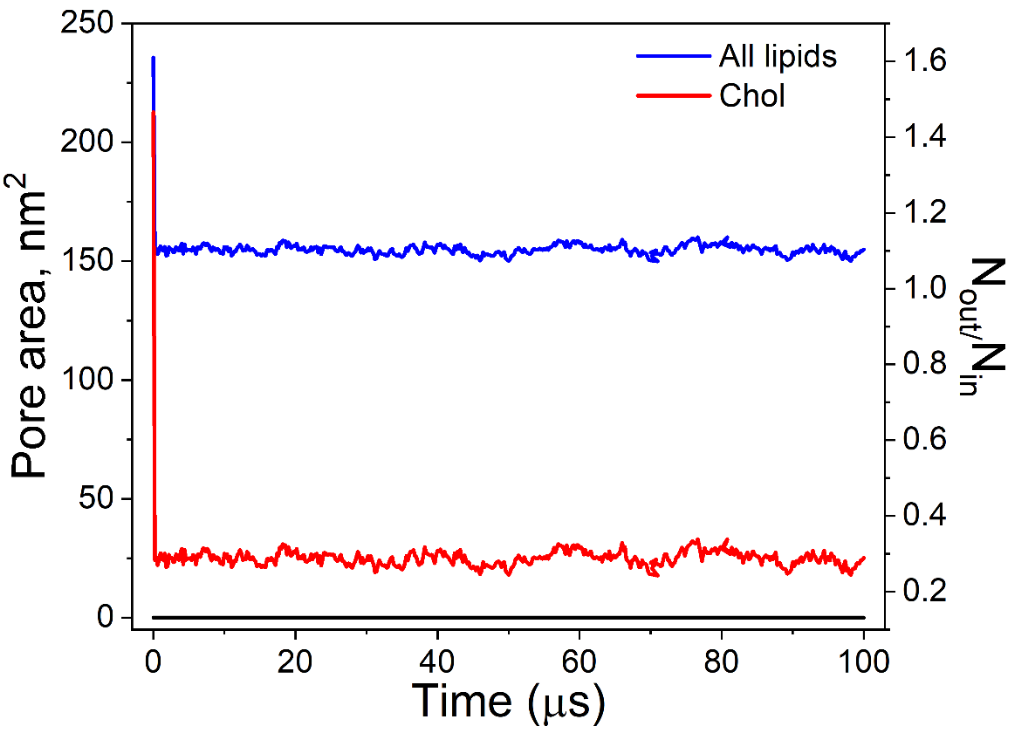


**Figure S7.** Lipids do not spontaneously undergo flip flop at long simulation times in intact membranes. The data shows the ratio of all lipids (blue) and chol (red) in the outer/inner leaflets of a DOPC:DOPE:DOPG:Chol bilayer patch exhibiting an initial 40% number asymmetry for a constrained bilayer. Note that chol immediately spontaneously flips to the inner leaflet ($N_{out}/N_{in} < 1.4$).

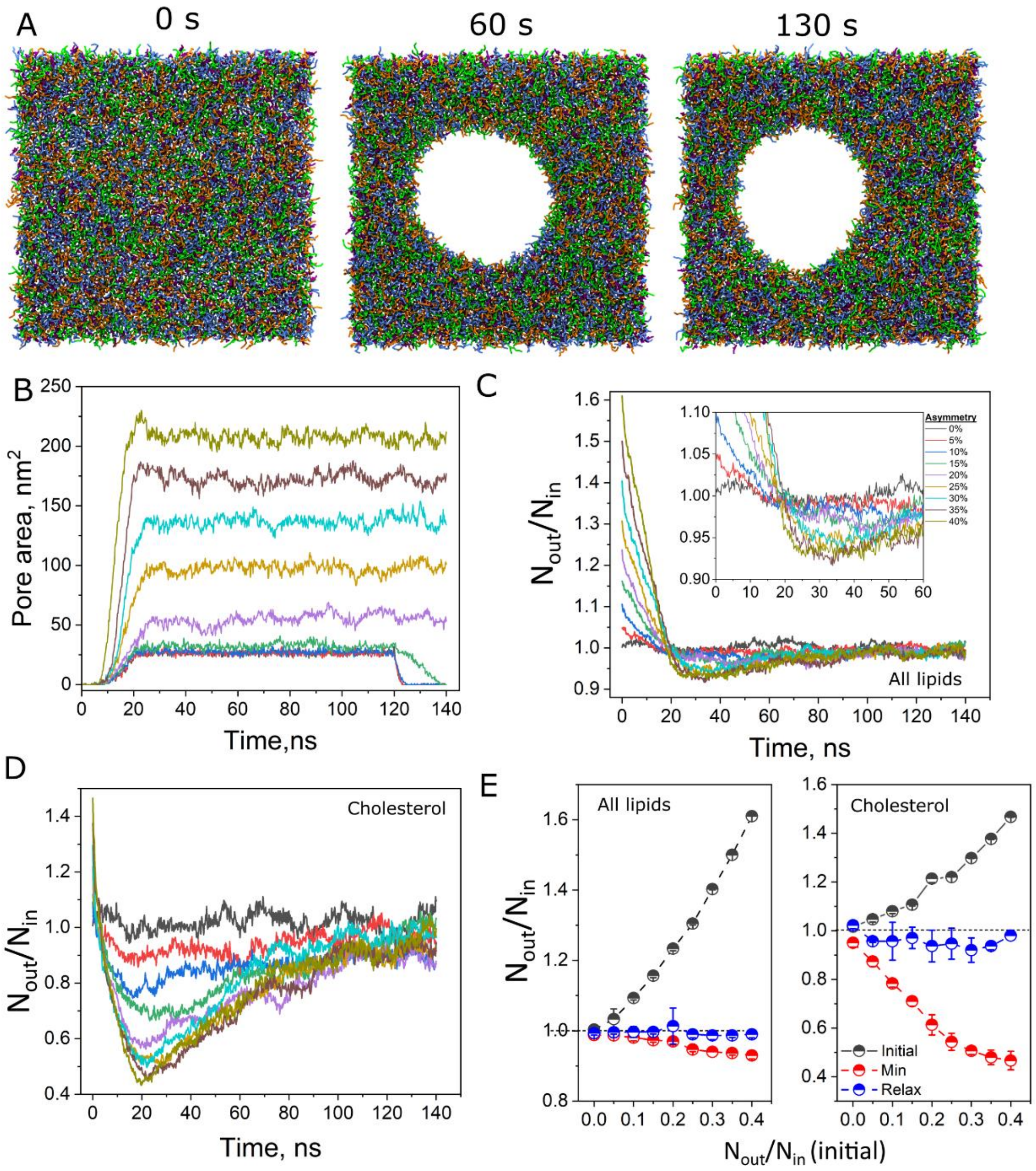


**Figure S8.** Coarse grained molecular dynamic simulations of asymmetric constrained membranes. Representative snapshots of a MD simulation of a DOPC:DOPE:DOPG:Chol (equimolar fractions) membrane patch exhibiting 40% number asymmetries at different (indicated) timepoints. The membrane is allowed to relax in all directions (laterally compressible membranes). C, dynamic evolution of pore area for different initial transbilayer asymmetries. C-D, lipid number asymmetry ($N_{out}/N_{in}$) for all lipids (Figure C) and for cholesterol only (Figure D). Inset in C: zoom-in in the early stages of pore opening. E, $N_{out}/N_{in}$ as a function of initial (pre-defined) asymmetries for all four lipids (left) and for cholesterol only (right) before lipid equilibration (initial, black data), upon full equilibration and before pore opening (min, red data) and after relaxation upon pore opening-closure (relax, blue data). Each data point represents the mean and s.d. from three independent simulations.

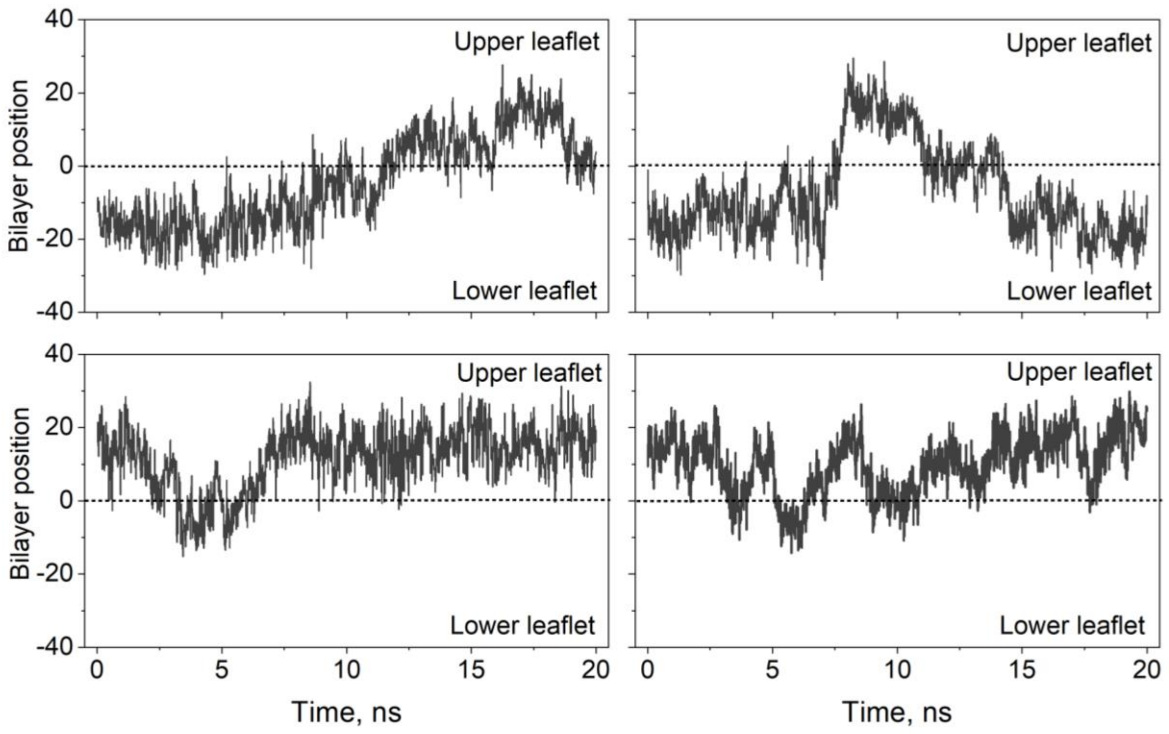


**Figure S9.** Single cholesterol flip flop in intact membranes and symmetrical membranes. Representative dynamics of four individual cholesterol molecules undergoing flip flop. Upper and lower leaflets are indicated.

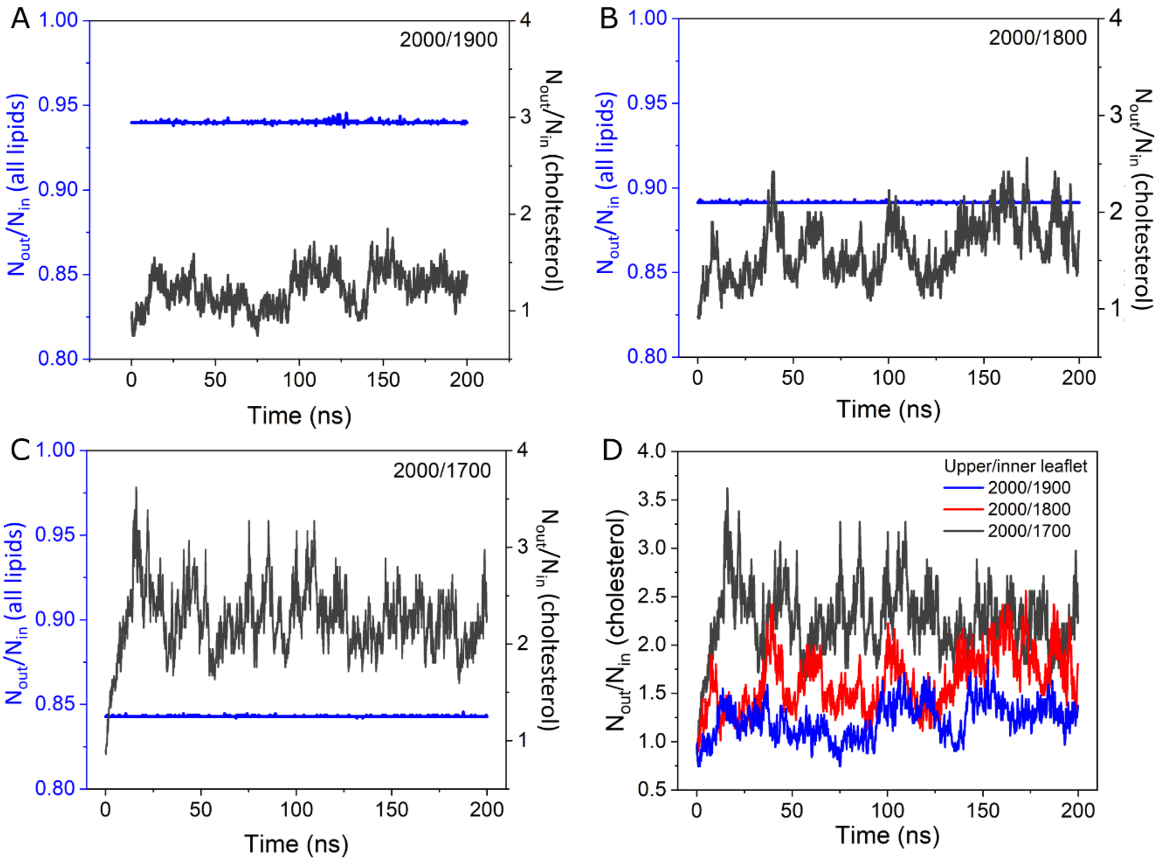

**Figure S10.** Cholesterol, but not phospholipids, undergo flip flop in asymmetric membranes. A-C, Asymmetry ($N_{out}/N_{in}$) of all lipids (blue) and cholesterol (black) of an intact and laterally constrained membrane patch with increasing levels of asymmetry. Lipid number in the outer/inner leaflets are indicated. D, cholesterol dynamics for the three asymmetries shown in A-C.

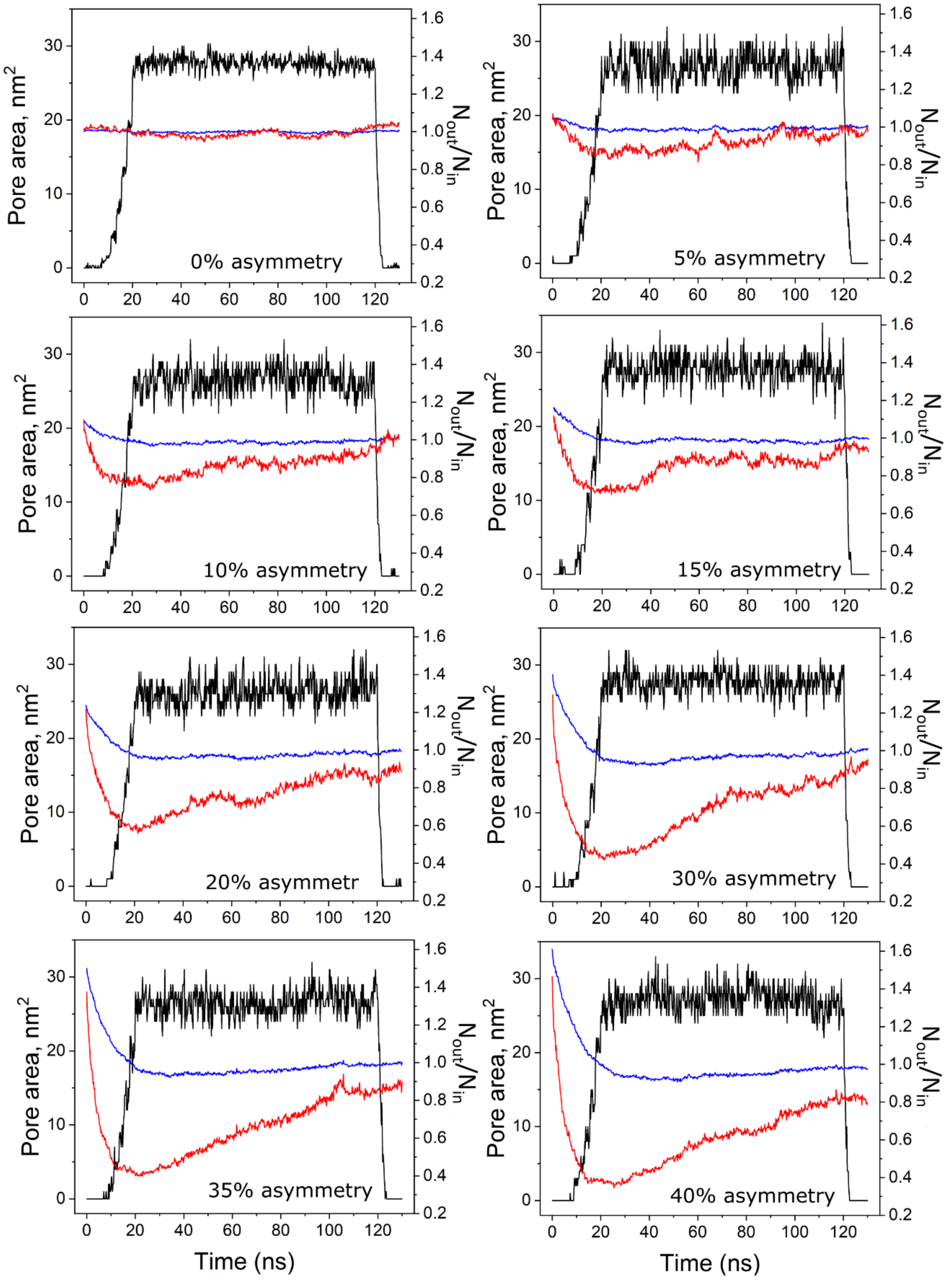


**Figure S11.** Asymmetry relieving upon pore opening in laterally compressible membrane. Black data: pore opening/closure. Lipid number asymmetry ($N_{out}/N_{in}$) for all lipids (blue) and cholesterol (red). Each plot represents the increasing initially imposed asymmetry.

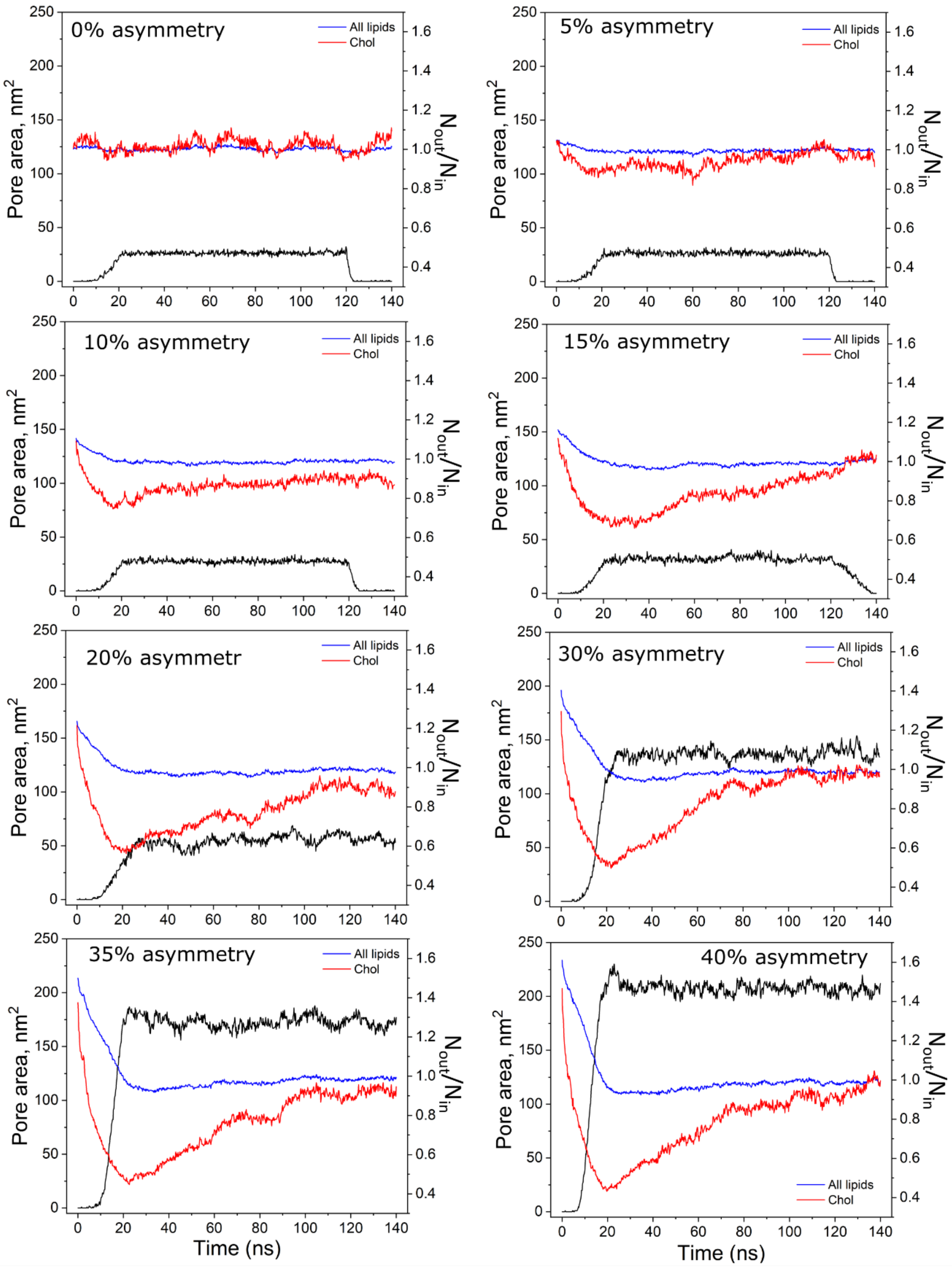


**Figure S12.** Asymmetry relieving upon pore opening in laterally constrained membrane. Black data: pore opening/closure. Lipid number asymmetry ($N_{out}/N_{in}$) for all lipids (blue) and cholesterol (red). Each plot represents the increasing initially imposed asymmetry.

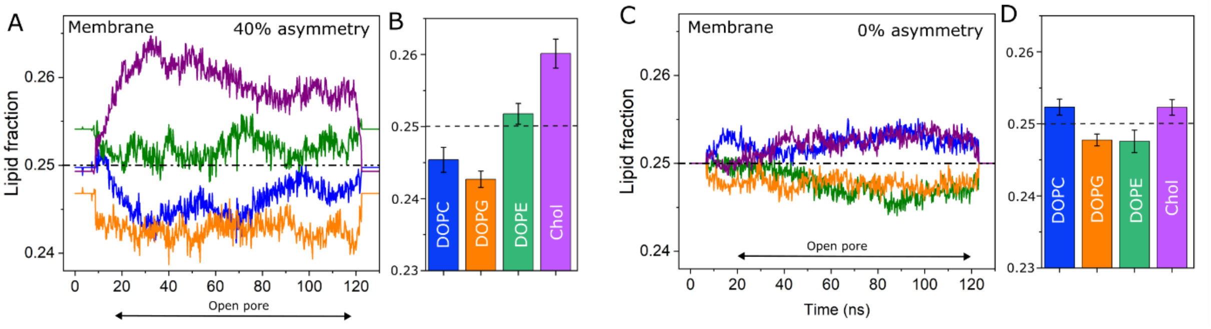


**Figure S13.** Lipid sorting in regions in intact membranes due to pore formation. A,C: dynamic fraction of lipids as a function of time 5 nm away from the pore rim at 40% (in A), and 0% asymmetry (in C). The time window where the pore is open (t = 20–120 ns) is indicated. B and D, average fraction of lipids in intact membranes for 40 % (in B), and 0% asymmetry (in D). The horizontal hatched lines at 0.25 represents lipids' expected distribution in the absence of compositional preferences.

**Supporting Videos**

**Video S1:** Pore closure of an electroporated control (no liposomes) DOPC:DOPE:DOPG:CHOL (equimolar fractions) GUV. Pulse strength 3 kV/cm and 15-200 μs pulse duration The time is indicated. GUV diameter: ~ 40 μm.

**Video S2:** Bursting of an electroporated DOPC:DOPE:DOPG:CHOL (equimolar fractions) GUV post incubation with 10 μM (lipid concentration) DOTAP:DOPE (50:50 molar fraction) LUVs. Pulse strength 3 kV/cm and 15-200 μs pulse duration The time is indicated. GUV diameter: ~ 25 μm.

**Video S3:** Control living HEK cells do not internalize water soluble probes. The cells were incubated with SRB (red, 10 μM) and Dextran 10 kDa (0.1 mg/ml) for 30 minutes. Incubation was carried out at 37 °C.

**Video S4:** Living HEK cells incubated with 500 μM (total lipid concentration) DOTAP:DOPE liposomes (50:50 mole fraction) become permeable to the small probe SRB. Note that strong dye signal in the cell cytosol. The cells were incubated with SRB (red, 10 μM) and Dextran 10 kDa (0.1 mg/ml) for 30 minutes. Incubation was carried out at 37 °C.

**Video S5:** 100 ns simulation of a laterally constrained membrane patch composed of DTPC (short tailed) phospholipids with 14% lipid number asymmetry.

**Video S6:** 130 ns simulation of a compressible DOPC:DOPE:DOPG:CHOL membrane patch. From 10 ns to 120 ns a pore is artificially opened.

**Video S7:** 130 ns simulation of a compressible DOPC:DOPE:DOPG:CHOL membrane patch with 40% lipid number asymmetry. From 10 ns to 120 ns a pore is artificially opened.

**Video S8:** 130 ns simulation of a laterally constrained DOPC:DOPE:DOPG:CHOL membrane patch. From 10 ns to 120 ns a pore is artificially opened.

**Video S9:** 130 ns simulation of a laterally constrained DOPC:DOPE:DOPG:CHOL membrane patch with 40% lipid number asymmetry. From 10 ns to 120 ns a pore is artificially opened.

**Video S10:** Cholesterol flipflop in an intact DOPC:DOPE:DOPG:CHOL membrane. Cholesterol is highlighted in yellow, the other lipids are indicated in blue.

**Video S11:** Lateral view of the pore upon pore formation over the course of 35 ns, showing lipid flip flop. One representative molecule of each phospholipid is highlighted, whose flipping time is approximately 20 ns. For the remainder of the membrane, only the phosphate groups are shown.

**Video S12:** Lateral view of a laterally constrained DOPC:DOPE:DOPG:CHOL membrane exhibiting 40% number asymmetry over the course of 130 ns (full length of the simulation), showing cholesterol flip flop. The cholesterol molecules are depicted in green/orange. For the remainder of the membrane, only the phosphate groups are shown in grey.

**References**

1. Lira, R. B. *et al.* Fluorescence lifetime imaging microscopy of flexible and rigid dyes probes the biophysical properties of synthetic and biological membranes. *Biophys. J.* **123**, 1592–1609 (2024).
